# Complete chromosome 21 centromere sequencing of families with Down syndrome reveals centromere size asymmetry

**DOI:** 10.1101/2024.02.25.581464

**Authors:** F. Kumara Mastrorosa, Alessia Daponte, Julie Wertz, Allison N. Rozanski, William T. Harvey, David Porubsky, Jordan Knuth, Gage H. Garcia, Marcelo Ayllon, Katherine M. Munson, Kendra Hoekzema, Weiya He, Stephanie L. Sherman, Emily G. Allen, Tracie C. Rosser, Claudia Rita Catacchio, Mario Ventura, Glennis A. Logsdon, Evan E. Eichler

## Abstract

Down syndrome, the most common form of human intellectual disability, is caused by nondisjunction and chromosome 21 trisomy (T21). Small centromeres have been hypothesized to contribute to its aetiology and studies on mammals suggest that larger centromeres are more efficiently transmitted, yet complete sequencing of chromosome 21 (chr21) centromeres has been particularly challenging. Using long-read sequencing, we sequenced and assembled the centromeres from eight families that include a child with free T21 (1 trio, 6 child–mother duos, and 1 singleton) all resulting from maternal meiosis I errors. Two of these families carry the smallest chr21 centromeres (143 and 181 kbp) observed in female individuals to date, exhibiting a ∼10.7- and ∼19.4-fold centromeric α-satellite higher-order repeat array size difference between the maternally inherited homologs, respectively. In both cases, the longer centromere harbors a poorly defined centromere dip region, marked by DNA hypomethylation, in the proband but not in the mother. A comparison of all proband chr21 centromeres (*n=*24) to those of controls (*n=*261) shows that small centromeres are not enriched in families with T21 (p-value=0.73); contrarily, chr21 extreme centromere size asymmetry (>10-fold) is unique of T21 (p-value=0.003), suggesting that this feature may represent a genetic risk factor for a subset of families with free T21. Additionally, phylogenetic reconstruction reveals that human chr21 has been particularly prone to such variation with some of the biggest size differences occurring over the last ∼17 thousand years of human evolution.

## INTRODUCTION

Trisomy 21 (T21) is the most common autosomal chromosomal aneuploidy in humans^1^ and the most common genomic cause of intellectual disability^2^. The clinical condition, known as Down syndrome, in most cases (>95%) is caused by the presence of an entire extra chromosome 21 (chr21) in the cells of the individual with the syndrome (free T21)^2^. Other cases are caused by a portion or the entire long arm of chr21 translocated to another chromosome as a result of Robertsonian translocations t(14;21) or t(21;21) or mosaic occurrences^2^. The majority of cases arise as a result of maternal meiosis I errors (MMIEs; ∼66%), followed by maternal meiosis II errors (MMIIEs; ∼21%), paternal meiosis I (∼3%) and meiosis II (∼5%) errors, or mitotic events after the zygote formation (∼5%)^2^.

Centromeres play an essential role in accurate chromosome segregation during meiosis by promoting equal segregation of homologous chromosomes and sister chromatids. Human centromeric regions consist of different types of satellite DNA, with the most abundant type being α-satellite: a ∼171 base pair (bp) nearly identical sequence tandemly repeated up to several megabase pairs (Mbp) in length^3^. For most human chromosomes, the functional centromere is composed of α-satellite repeats organized into higher-order repeat (HOR) arrays that tend to diverge in sequence at the pericentromeric edges. Until recently, the complete sequence of all human centromeres was unknown because they could not be assembled with standard short-read sequencing data. However, in the last five years, improvements in long-read sequencing technologies and computational assembly methods have made it possible to assemble centromeres^4–7^, the short arm of acrocentric chromosomes^8,9^, large segmental duplications^10,11^, and other complex loci^8,12,13^ previously regarded as inaccessible.

Functionally, the kinetochore is marked by the presence of nucleosomes containing the histone variant H3 CENP-A, which provides the foundation for the assembly of inner- and outer-kinetochore proteins and plays a crucial role in cell division and chromosome segregation^14,15^. The analysis of the complete sequence of human centromeres^5–7^ has made it possible to define the likely site of kinetochore attachment: a pocket of hypomethylated CpG DNA called the centromere dip region (CDR)^16,17^, where chromatin immunoprecipitation followed by sequencing (ChIP-seq) shows an enrichment of CENP-A. All completely sequenced human centromeres to date derive from primary tissue (blood) or cell lines and possess CDRs up to tens of kilobase pairs (kbp) in length where CENP-A accumulates, implicating this as a general epigenetic signature of the kinetochore^7^.

Given their role in cell division, centromeres, and in particular length differences, have been postulated to play an important role in chromosome nondisjunction, including Down syndrome^18,19^. Some studies on mice suggest that centromeres with longer repeat arrays are preferentially retained in the egg during maternal meiosis I^20–23^ and that in general, centromeric features, such as sequence and length, and kinetochore components^23–26^ are correlated: a concept known as “centromere strength.” A recent investigation in humans also suggested that individuals with T21 might uniformly have smaller chr21 centromeres (possibly as a result of reduced size of the centromeric α-satellite HOR array)^27^. Centromere size was estimated, however, based on quantitative, real-time PCR^27^, which can be challenging given the high degree of sequence identity between chromosome 13 and 21 α-satellite DNA and a limited understanding of the sequence diversity within the allelic α-satellite HOR array^27^. Other studies showed that cohesion loss between sister centromeres^28^ and reduced recombination^29^ increase the risk of nondisjunction. Nevertheless, the most well-established nongenetic risk factor is increased maternal age at conception^19^, though there is not a well-established genetic risk factor for free T21.

It is currently unknown whether centromere variation amongst humans increases the likelihood of nondisjunction. For instance, centromere size, sequence, allelic differences, and epigenetic features might influence nondisjunction risk and indicate predisposition to such events.

Recently, combining both Pacific Biosciences (PacBio) high-fidelity (HiFi) long-read sequencing and ultra-long Oxford Nanopore Technologies (UL-ONT) sequencing, we demonstrated that it is possible to completely sequence and distinguish human autosomal alleles for chr21^7^. Here, we present the first complete linear sequence and epigenetic profile of chr21 centromeres in eight individuals with MMIE free T21 and eight of their parents (7 mothers and 1 father). We also compare genetic and epigenetic features of the chr21 centromeres to a control group of 261 completely sequenced chr21 centromeres from two hydatidiform moles and control individuals generated as part of the Human Genome Structural Variation Consortium^30^ (HGSVC) and Human Pangenome Reference Consortium^31^ (HPRC) to reveal unique features of T21 centromeres.

## RESULTS

### Maternally inherited chr21 centromeres show size asymmetry in an individual with T21

To study the chr21 centromeres of an individual with T21, we initially sequenced the DNA of a proband (maternal age at childbirth = 29) using both PacBio HiFi sequence data (59×) and UL-ONT sequence data (23×). We used Verkko, a hybrid genome assembly workflow^32^, to assemble all proband chr21 centromere haplotypes (total genome assembly size = 6.118 Gbp, contigs = 3,466, contig N50 = 7.72 Mbp). After standard validation and quality control^5,7^ (**Methods**, **Supplementary Figure 1**), sequence assembly of phased chr21 centromere haplotypes confirmed an MMIE where haplotypes 1 (H1) and 2 (H2) arose from maternal nondisjunction and haplotype 3 (H3) was inherited from the father (**Figure 1a**).

**Figure 1.**
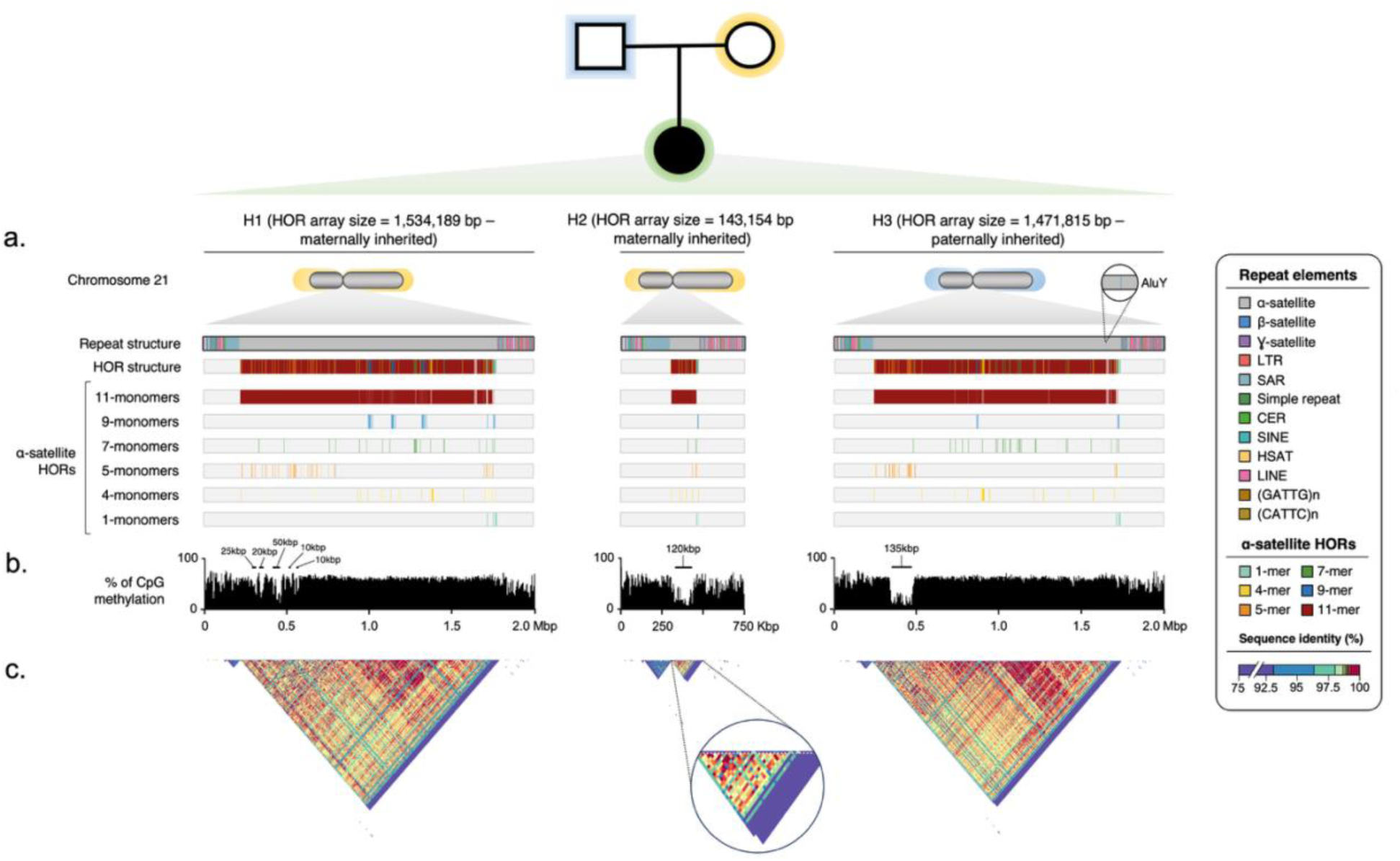
Genetic composition and epigenetic profile of three chromosome 21 (chr21) centromeres from an individual with Trisomy 21 (T21). **a)** Size and repeat composition of three centromeric haplotypes for an individual with T21 (Coriell ID: AG16777). Different α-satellite higher-order repeat (HOR) monomers mapping within the α-satellite array are colored (see legend). Only α-satellite HOR monomers present more than 10 times in at least one haplotype are shown (see **Supplementary Table 1** for complete monomer composition). **b)** Methylation profile of each centromere with hypomethylated pockets indicating the location of the kinetochore. **c)** Pairwise sequence identity heatmaps (StainedGlass^33^). Sequence identity is depicted in 5 kbp windows across the centromeres revealing regions of high- and low-identity homology within each α-satellite HOR array (see legend).

H1 and H3 centromeres are the most similar in size with an α-satellite HOR array of ∼1.53 Mbp and ∼1.47 Mbp, respectively. By contrast, maternal H2 is 10.7-fold smaller, with an α-satellite HOR array of only 143 kbp (**Figure 1a**). The α-satellite HOR array sequences from all three haplotypes are predominantly composed of 11-monomer α-satellite HORs (∼86-93%) with a smattering of rarer α-satellite HORs 2-, 10-, 13-, 17-, and 18-monomers in length (each constituting <3% of the sequence) (**Figure 1a; Supplementary Table 1**). Other than their size, the three chr21 centromeres share a remarkably similar organization, with the α-satellite HOR array flanked by human satellite 1a (HSat1a; annotated as “SAR’’ by RepeatMasker) on the p-arm and various classes of mobile elements (SINEs and LINEs) on the q-arm. Also, H3 is distinguished by a 317 bp AluY retrotransposon insertion in its α-satellite HOR array (**Figure 1a**).

Given the use of long-read sequencing, which allows for CpG methylation to be detected, we were able to assess the position and distribution of each chr21 centromere CDR, the pocket of hypomethylated DNA indicating the location of the kinetochore^5,6,17^. Both H2 and H3 match this description with a CDR of 120 kbp and 135 kbp, respectively. By contrast, the maternally inherited H1 does not have a clearly defined CDR. The CDR is fragmented into five smaller pockets ranging from 10 to 50 kbp and spanning across 280 kbp of centromeric α-satellite HORs (**Figure 1b**). For each CDR, we calculated the percentage of methylated CpGs (defined as having ≥25% methylated CpGs at the same position; **Methods**) across the region comprising the CDR pockets (from the start of the first CDR window to the end of the last one). We found that the percentage of methylated CpGs within H1’s CDR was much higher than those of H2 and H3 CDRs (∼71% vs. ∼39% and ∼38%, respectively; **Figure 1b**). This more diffuse pattern of CDRs is associated with the largest HOR array within this family. We orthogonally validated the centromere methylation profiles using PacBio HiFi sequencing data (**Supplementary Figure 2**).

Because of the central role of centromeric proteins, such as CENP-A and CENP-C, in defining the site of kinetochore assembly other than the CDRs, we assessed the level of CENP-A and CENP-C on chr21 centromeres in proband AG16777 and their parents using an immunofluorescence (IF) combined with fluorescence *in situ* hybridization (FISH) targeting the α-satellite sequence, CENP-A, and CENP-C proteins (**Supplementary Figure 3**). We observed, however, no statistically significant differences in CENP-A or CENP-C signal intensities in any member of the family.

For each chr21 centromere, we also generated sequence identity heatmaps, based on pairwise sequence alignment of 5 kbp segments using StainedGlass^33^ (**Figure 1c**). All three haplotypes show similar compositional features, with the most identical segments mapping to the central section of the α-satellite HOR array but not corresponding to the CDR, with the exception of H2 due to its smaller size. This structure is also similar to the complete chr21 α-satellite HOR arrays described for two complete hydatidiform moles (CHM1 and CHM13)^5,6^, as well as population samples^7^, and generally follows the Smith organizational model with monomeric satellite at the edges as previously described in 1976^34^. For both H1 and H3, the CDR and the predicted site of kinetochore attachment occur peripherally in a region composed of more heterogeneous α-satellite HOR cassettes (with an admixture of 5- and 11-monomer HORs).

For this index family, we also studied additional proband centromeres (chromosomes 4, 12, 15) for which both haplotypes were correctly assembled (**Supplementary Figure 4**). Chromosome 12 homologs are the only ones having the CDRs roughly in the same relative position on both homologs. Chromosome 4 and 15 haplotype CDRs are positioned differently instead and one of chromosome 4 is divided into three windows intervaled by small highly methylated regions (**Supplementary Figure 4**) confirming the CDR heterogeneity recently observed by Logsdon et al. from their analysis of cell line samples as part of the HGSVC^35^.

### Parental chr21 centromeres show transgenerational methylation changes and suggest a preferential CDR position

To understand the inheritance of the chr21 centromeres, we sequenced and assembled the genomes of both parents using DNA extracted from lymphoblastoid cell lines and fully reconstructed each of the centromere haplotypes from the diploid parental cell lines. We generated PacBio HiFi sequencing data at 37× coverage for the mother and 40× for the father and complemented these data with UL-ONT data at 25× and 12× coverage, respectively, to generate *de novo* assemblies (mother: total genome assembly size = 6.078 Gbp, contigs = 4,827, contig N50 = 5.41 Mbp; father: total genome assembly size = 5.947 Gbp, contigs = 5,032, contig N50 = 5.01 Mbp). Centromeric contigs from chr21 were extracted, validated, and compared to the child with T21 (see **Methods** for assembly correction).

Sequencing confirmed that the largest and the smallest chr21 centromeres (H1 and H2, respectively) were maternally transmitted, while H3 originated from the father (Figure 2a-b). Sequence identity between parent and proband haplotypes was >99.99% identical, confirming the accuracy of the assembly as well as the stability of the centromere transgenerationally and in cell culture (Supplementary Figure 5). No difference in the α-satellite repeat composition, organization, or array size was observed, except for a 1 bp deletion in the poly(T) tract associated with an AluY insertion on the paternal haplotype (H3) when compared to the proband (Supplementary Figure 6). We note that the second non-transmitted paternal haplotype (father H4) has an α-satellite HOR array size of 254 kbp (Figure 2a). This is 112 kbp larger than the small maternal haplotype (H2). Like the other haplotypes, the 11-monomer α-satellite HORs constitute the majority of its array sequence (∼86%), with 2- to 5-, 7-, 9-, 10-, and 13-monomer HORs at lower frequency (each composing <6% of the array sequence).

**Figure 2.**
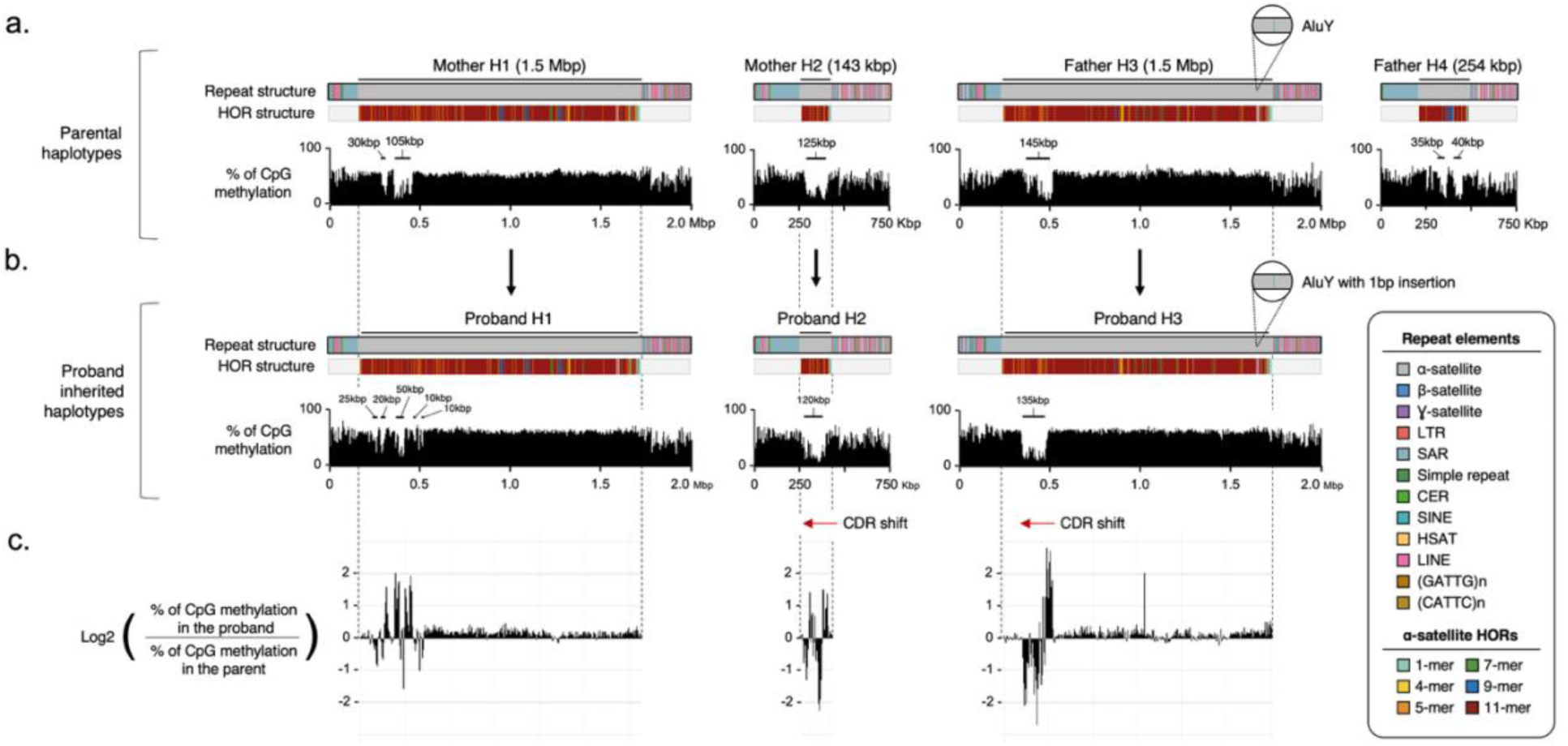
Genetic and epigenetic features of parent–proband transmission of chr21 centromeres. Composition and organization of α-satellite HOR arrays (only α-satellite HORs present more than 10 times in at least one haplotype are shown; see **Supplementary Table 1** for complete HOR composition), with CpG methylation profiles of **a)** parental chr21 centromeres and **b)** transmitted proband chr21 centromeres. **c)** Log_2_ ratio of CpG methylation frequency between proband and parental haplotypes.

Comparing the centromeric CpG methylation profiles between the parents and the proband, we find that the atypical methylation pattern observed in proband H1 (five hypomethylated pockets 10-50 kbp in size) appears to be specific to the proband (Figure 2a-b; see Supplementary Figure 2 for orthogonal validation). The mother shows a more typical CpG methylation profile, with two clear CDRs of 30 and 105 kbp in length, and ∼46% methylated CpGs across the region comprising the CDR pockets. This is 25% less compared to the proband (∼71%), and it contrasts with the transmitted maternal H2 (42% vs. 39% in proband) and paternal H3 (48% vs. 38% in proband), which show more similar CDR patterns. The CDRs of chr21 H2 and H3 centromeres differ by only an estimated 5-10 kbp in size between the generations (Figure 2c). We also observe a shift of both proband H2 and H3 CDRs towards the p-arm relative to the parents despite nearly identical sequences. The maternal H2 CDR, for example, starts 37 kbp from the beginning of the α-satellite array, while in the proband, the hypomethylated pocket starts 11 kbp upstream. The paternal H3 has a CDR initiating 131 kbp from the beginning of the α-satellite HOR array, which is shifted 25 kbp towards the p-arm in the proband (Figure 2c).

### Analysis of additional families with Down syndrome reveals another case of asymmetric chr21 centromeres

To validate the findings from the index trio, we generated PacBio HiFi and UL-ONT sequence data from seven additional families: six mother-proband duos where MMIEs had been previously identified and one singleton where parental DNA was not available.

Importantly, unlike the original trio, in six of these families both blood and cell line DNA material were available for investigation. Because the maternal age of the index case was 29 years, we specifically selected mothers representing a younger age range (20-34 years) (**Supplementary Table 2**). Despite the well-known maternal age effect with T21, we note, however, that this age range still represents the majority of the cases in the extended collection from the Atlanta and National Down Syndrome Projects^36,37^ (maternal age at birth <35 for ∼54% of the cases; 2,171/3,985), with a mean maternal age at birth of 32.9 years old. We assembled their genomes with the same methods used for the original family (probands mean genome assembly size = 6.072 Gbp, # of contigs = 1,237, contig N50 = 45.84 Mbp; mothers mean genome assembly size = 6.064 Gbp, # of contigs = 2,217, contig N50 = 35.16) and analyzed their chr21 centromeric sequences. Of the proband centromeres, 95% (20/21) were completely assembled while 67% (8/12) of the maternal centromeric haplotypes were assembled without any detectable error.

In this extended cohort, we identified one additional family showing extreme centromere size differences similar to that described in the original trio with T21. Duo 7210079 shows a 19.4-fold chr21 centromere size difference between maternal homologs (original trio: 10.7-fold) with a small centromere α-satellite HOR array of only 181 kbp (original trio: 143 kbp) and a longer one of ∼3.5 Mbp (original trio: 1.5 Mbp). In this duo as well, we observed a *de novo* epigenetic change in the centromeric methylation profile, with the proband having a less-defined CDR compared to the maternal for the largest haplotype (Figure 3). Since the 7210079.00-H3 centromere was not assembled completely in the proband, we leveraged the maternal locus (assembled contiguously with few sequence errors) as a reference to align UL-ONT reads both from mother and child and compare methylation profiles and CDR structure. Specifically, the maternal CDR maps to a unique window of 150 kbp more proximally toward the p-arm, while in the proband, the CDR is divided into five smaller windows (size range 75-10 kbp) despite remaining in the same genomic area (Figure 3). Mother 7210079.20 had a similar age at childbirth (age: 27) as the original trio mother (age: 29).

**Figure 3.**
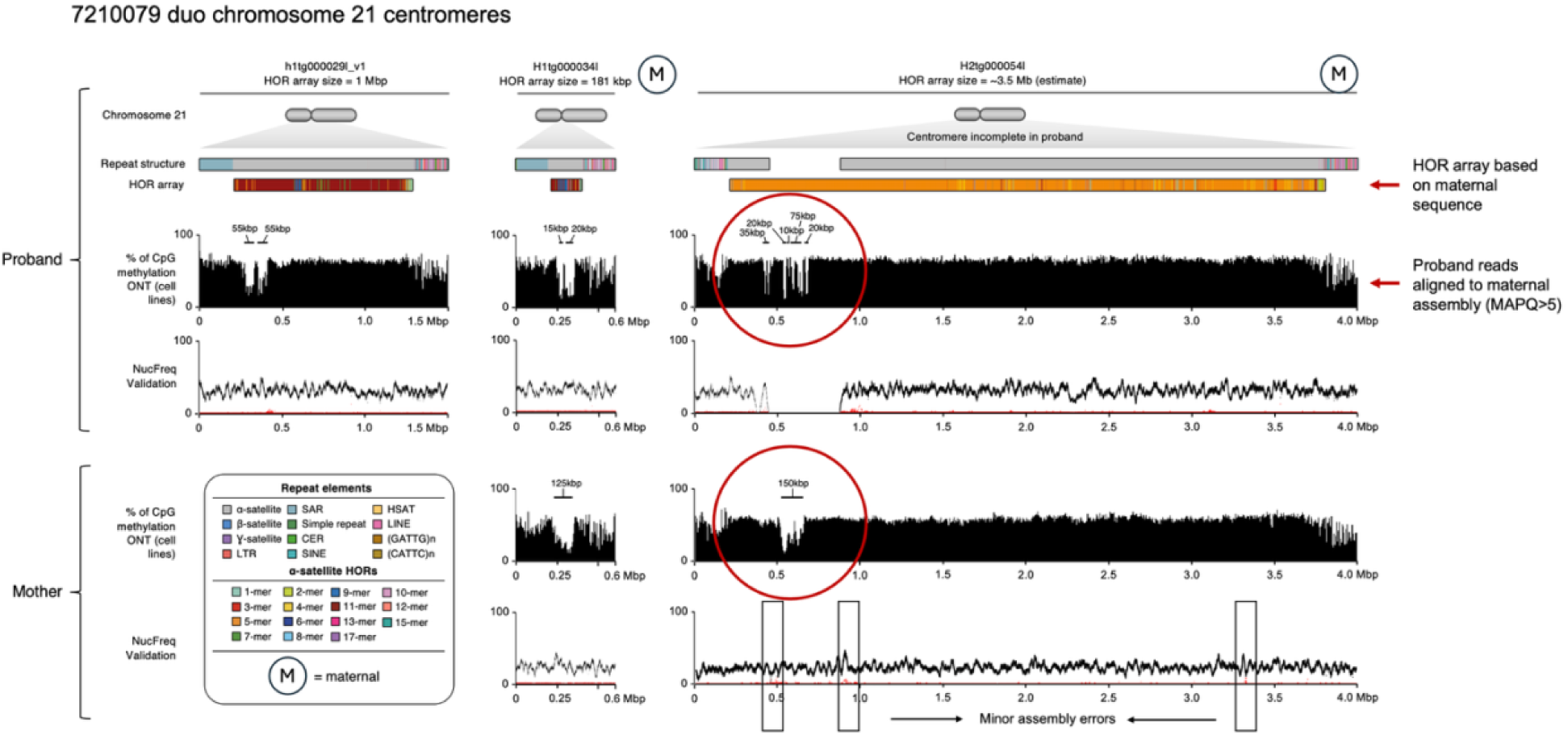
Chromosome 21 (chr21) centromeres in duo 7210079 exhibit size asymmetry and transgenerational methylation changes. Repeat structure, α-satellite HOR array organization, CpG methylation profile, and NucFreq validation of each chr21 centromere in 7210079 mother–child duo. Transgenerational DNA methylation changes are circled in red. The 7210079.00 proband H3 was not completely assembled. Since the same haplotype was assembled in the mother, we used the maternal assembly as reference to study the methylation profile in the proband.

A third family from the cohort with T21, 7210223, showed an epigenetic change between the transmitting parent and proband despite showing only a 2.6-fold centromere size difference between maternal homologs (587 kbp and 1.5 Mbp). Contrary to what we observed for the other two families with extreme maternal size difference, we observe a CDR shift occurring on the smaller maternally inherited centromere for proband 7210223.00. The 7210223.00-H2 centromere (size: 587 kbp) shows four CDR windows in the mother (size range 50-10 kbp) distributed across 170 kbp, while in the proband, we detect six smaller CDRs ranging in size from 10 to 20 kbp distributed across 310 kbp (Supplementary Figure 7). For all other families, only minor transgenerational changes in the CDR methylation profiles were observed for chr21 centromeres (Supplementary Figure 7). Among the remaining five probands, all have maternal homologs that are at least 1 Mbp large and a size difference of less than twofold (**Supplementary Table 2**).

With respect to sequence composition, most proband centromeric α-satellite HOR arrays in this cohort are similar to the original, composed predominantly of 11-monomer α-satellite HORs (∼77-94%), with rarer HORs representing the rest of the sequence. Notably, 4710291.00-H3 has 10% of its sequence composed of a 4-monomer α-satellite HOR, 7210239.00-H1 has 8%, and 7210008.00-H1 has 5%. The smallest haplotype of this cohort, 7210079.00-H1, has 9% of its sequence organized in a 9-monomer HOR (**Supplementary Table 1**). However, 7210079.00-H3 is unique compared to all the other centromeres. Based on the maternal α-satellite HOR array structure (Figure 3), it is the largest centromere sequenced amongst all characterized (α-satellite HOR array of ∼3.5 Mbp), with almost 1 Mbp more sequence than the second largest (4710291.00-H3). Also, its predominant α-satellite HOR array unit is composed of 5-monomer units (88%) with only 6.65% of its sequence made up of the more typical 11-monomer units.

Because both blood and cell line DNA were available for a subset of the probands, we compared the chr21 centromere methylation patterns for the two sample types for the five probands with complete centromeres. PacBio HiFi data were generated from blood DNA, and UL-ONT data were generated from cell line DNA. We confirm that the CDR positions remain the same from primary tissue (blood) to cell line (lymphoblastoid) (Supplementary Figure 8). This was concluded by observing ONT and HiFi data processed with the latest methylation callers dorado (v0.4.2) or guppy (6.3.7 or newer) and Jasmine (v2.2.1), respectively. We also confirmed the methylation callers’ concordance in the original trio AG167 (Supplementary Figure 2) and other samples (Supplementary Figure 8), for which the two long-read datasets were generated from cell lines. Despite a stable CDR position, we observed an overall higher level of methylation in blood and more readily defined CDR in lymphoblastoid cell lines (Supplementary Figure 8).

### Population chr21 centromere diversity and phylogeny

To understand genetic and epigenetic diversity in the general population, we initially selected 35 fully resolved chr21 centromere haplotypes from 16 different ethnic groups representing all five ancestry superpopulations as defined by the 1000 Genomes Project^38^. We observed considerable diversity in α-satellite HOR composition and length amongst these 35 complete chr21 centromeres. While the most prevalent α-satellite HOR is 11-monomers in length and comprises ∼83% to ∼95% of the α-satellite HOR array sequence, in most of the complete chr21 centromeres, we also detect less abundant α-satellite HORs that are 12, 14, 15, 16, and 20 α-satellite monomers in length (**Supplementary Table 1,** Figure 4a-b). There are two notable haplotypes whose composition differs markedly from the predominant 11-monomer α-satellite HOR organization: an individual of Gambian (GWD) descent, HG02647, carries a 4-monomer α-satellite HOR composing ∼15% of its H2 centromere sequence; and an individual of Punjabi (PJB) descent, HG03710, with 44% of its H1 centromere sequence composed of 5-monomer α-satellite HORs (Figure 4c**, Supplementary Table 1**). The 7210079.00-H3 described earlier is the only other chr21 centromere containing a majority of 5-monomer HORs (88%), and along with HG03710-H1, represent the two largest chr21 centromeres sequenced to date. Sequence identity plots (Figure 4c) also reveal that the 11-monomer-enriched α-satellite HOR region of HG03710-H1 shows greater sequence identity when compared to the 5-monomer-enriched regions. In contrast, the 4-monomer α-satellite HOR of HG02647-H2 shows greater sequence identity when compared to other flanking monomers (Figure 4c). If sequence divergence is a proxy for evolutionary age, our analyses may suggest that the 4-monomer α-satellite HOR has more recently evolved, while the 5-monomer α-satellite HOR is more ancient. Notwithstanding, both of these rarer centromere haplotypes map their CDRs to the noncanonical 5-monomer and 4-monomer α-satellite HORs.

**Figure 4.**
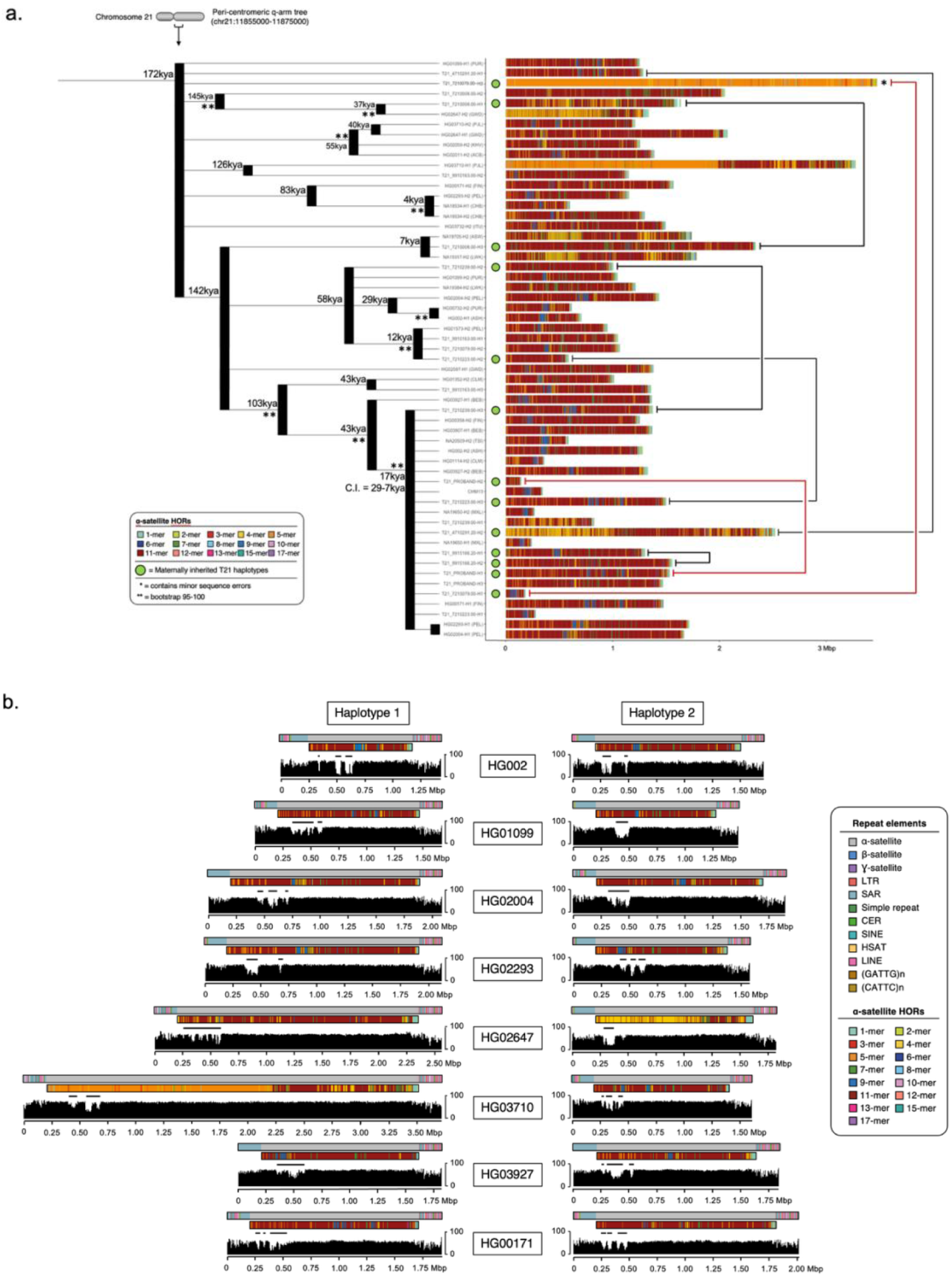
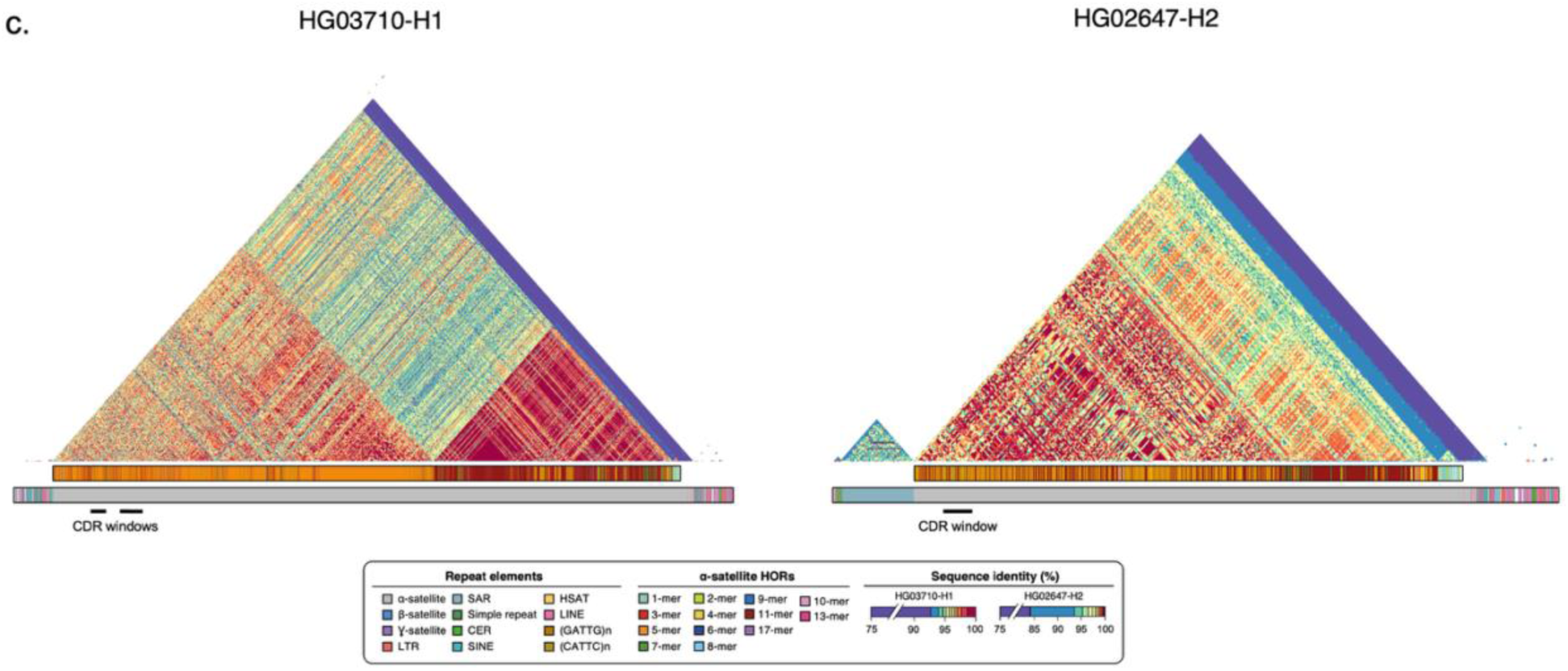
Diversity in α-satellite HOR array length, composition, and CpG methylation profile amongst diverse chr21 centromere haplotypes. **a)** Maximum-likelihood phylogenetic tree based on a multiple sequence alignment of a 20 kbp segment from the pericentromeric q-arm regions from 57 unique completely sequenced chr21 centromeres. The HOR array structure of each is depicted and estimated divergence time for each cluster is projected onto the nodes. Bootstrap values 95 to 100 (**) are marked with asterisks. Black lines connect homologous chr21 centromeres found in mothers, which were both transmitted to their child with T21. Red lines indicate the homologs exhibiting extreme centromere size asymmetry. **b)** CpG methylation profiles of 16 population haplotypes (see also Supplementary Figure 9). **c)** Pairwise sequence identity heatmaps (generated with StainedGlass^33^ using 5 kbp windows) across chr21 centromeres enriched with 5-monomer and 4-monomer α-satellite HORs (see legend).

We also compared the diversity in length and location of the CDRs for the various chr21 centromere haplotypes. The number of CDRs typically range from 1 to 4 and span a total of 90-335 kbp in total length (Figure 4b**; Supplementary** Figure 9). None show patterns similar to the one observed in the proband (H1) with multiple small CDRs mapped in close proximity.

However, some haplotypes, such as HG02647-H1 and HG03927-H1, show large CDR windows with peaks of higher methylation within. We observe four chr21 centromere haplotypes that have two hypomethylated regions separated by more than 100 kbp: HG002-H1 and H2; HG02293-H1; and HG01573-H2. The dispersed CDRs tend to be smaller (10 to 25 kbp) and contained within a single pocket, except for HG01573-H2, which has two. Irrespective of sequence composition, 91% (20/22) of the chr21 CDRs map to the p-arm of the centromere, suggesting a polarity or preference for CDR location (Supplementary Figure 9).

We also constructed a chr21 centromere phylogenetic tree using 20 kbp of flanking pericentromeric sequence in the q-arm from the genomes assembled in this study and chimpanzee as an outgroup (**Methods**). We have shown previously that these flanking sequences can serve as a reasonable proxy of evolutionary history due to the large block of linkage disequilibrium created by the suppressed recombination around the centromere^7^. We find that 50% (11/22) of T21 haplotypes included in the analysis are part of a large monophyletic clade comprising 40% (23/57) of all chr21 centromeres (Figure 4a). The clade harbors the largest (H1) and smallest (H2) centromeres for proband AG16777, which are predicted to have arisen from a common ancestor 17 thousand years ago (kya), despite the fact that these two haplotypes vary by more than 10-fold in their overall length. It also contains the small centromere found in the second family (7910079) showing extreme centromere size asymmetry. We note that this particular clade is significantly enriched for centromeres smaller than 500 kbp (Fisher’s exact test p-value = 0.001). In contrast, our analysis also reveals that the two large chr21 centromeres containing an abundance of 5-monomers α-satellite HORs (7210079.00-H3 and HG03710-H1) diverged much earlier (126 and 172 kya) compared to the other haplotypes. The low degree of sequence identity with other canonical 11-monomers (Figure 4a) confirms an ancient origin, possibly the result of an archaic introgression.

### Maternal asymmetry in T21 centromeric α-satellite HOR length

To establish a baseline for centromere length distributions in the human population, we compared the T21 centromeres to an expanded set of 261 completely assembled chr21 centromeres from control individuals. The control cohort consists of two hydatidiform mole (CHM13 and CHM1) chr21 centromeres, 184 chr21 centromeres (92 samples) from the recently released genomes part of the HPRC^31^ (year 2 release), 40 chr21 centromeres recently published by the HGSVC^30^, and an additional 35 from the two consortia characterized with the same methods applied to the families with T21 (**Supplementary Table 3**). The centromeres from the HPRC year 2 release were characterized with CenMAP (v0.3.1), a recently released centromere mapping and annotation pipeline (https://github.com/logsdon-lab/CenMAP). Comparing the size distribution of all control haplotypes (*n*=261) to those from individuals with T21 (*n*=24), we find no statistically significant difference (t-test p-value = 0.92) (Figure 5a).

**Figure 5.**
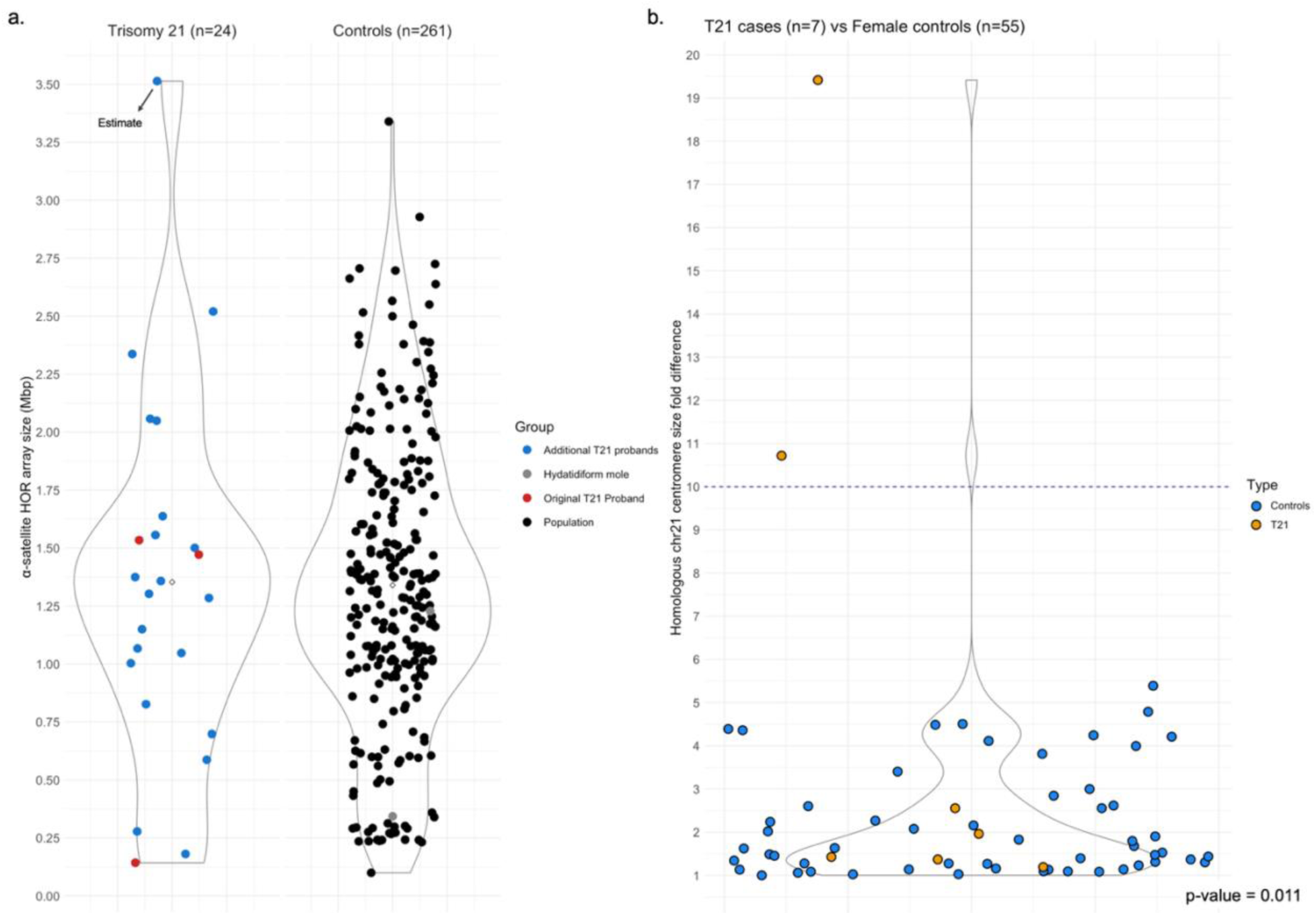
Comparison between centromere HOR array sizes from individuals with T21 and controls. **a)** Size distribution comparison of chr21 centromeric α-satellite HOR arrays from individuals with T21 and population samples. This plot includes the HOR array sizes of 24 T21, 2 hydatidiform moles, and 259 population haplotypes. **b)** Homologous chr21 centromere size fold differences in individuals with T21 (only maternally inherited centromeres; orange) and female population samples (blue). P-value indicates a significant difference in extreme asymmetry cases between the two groups.

Similarly, the frequency of small centromeres (<500 kbp) also showed no significant difference between the two groups (3/24 in T21 and 27/261 in the general population groups; Fisher’s exact test p-value = 0.73) despite an earlier report suggesting the contrary^27^. Notwithstanding, the small centromeres identified in AG16777 and 7210079 families (mothers and children) remain the smallest chr21 α-satellite HOR array sequenced to date in any female individual, measuring 143 kbp and 181 kbp respectively (Figure 5a). For the male individuals, only one centromere was found to have a smaller α-satellite HOR array: HG00739-H1, which represents the smallest chr21 centromere ever sequenced with an array of only 100 kbp.

Next, we considered the diplotype of the transmitting mothers and assayed the frequency of extreme centromere size asymmetry (>10-fold) in the population using 116 control samples (females = 55; males = 61) and seven individuals with T21 for which both centromere haplotypes were fully resolved (the singleton with T21 was excluded since we could not phase the centromeres nor determine if the centromere size remained the same between the two generations; Supplementary Figure 10). A significant difference in extreme homologous chr21 centromere size asymmetry cases was observed between mothers with children with T21 and the general population. We identified two cases in the cohort of individuals with T21 and none in 116 control individuals where homologous centromere HOR length differed more than 10-fold (Fisher’s exact test p-value = 0.003) (Supplementary Figure 10). Given the fundamental differences in male and female meiosis, we repeated the analysis restricting further to only female individuals from the general population. The number of asymmetrical centromeres remained statistically significant for families with T21 (Fisher’s exact test p-value = 0.011) (Figure 5b). We further expanded the controls by including another 23 female and 22 male individuals from the HPRC year 2 release that were completely assembled but had small errors in their sequences (e.g., collapses, misjoins, etc.). In these samples, the centromere size estimate is not as accurate, but we do not expect a great difference between the estimate and their actual sizes. Once again, none exhibited extreme centromere asymmetry. If we consider these additional female samples in our comparison, the difference becomes even more statistically significant (Fisher’s exact test p-value = 0.006).

## DISCUSSION

Down syndrome was first described in 1866^39^ and its typical T21 karyotype description was made in 1959^40^. Despite extensive research over the last 65 years, very little is still known about the genetic etiology of this condition. To this day, we still do not have a method to assess predisposition to this condition nor are aware of any direct genetic risk factor that leads to free T21. We could only test directly for T21 during pregnancies with invasive procedures, such as amniocentesis until more recent, safer noninvasive prenatal tests emerged^41^. Understanding the genetic architecture of T21 and genetic factors that predispose to it may be another important consideration in its prevention.

In this study, we present for the first time the complete sequence and epigenetic characterization of chr21 centromeres of eight individuals with free T21, seven of their mothers, and one father. In all families, T21 was caused by MMIEs and all mothers were <35 years old at childbirth. We initially aimed to investigate chr21 centromere variability in individuals with T21 and test the hypothesis that individuals with the syndrome carry small centromeres. Previous studies postulated that “centromere strength” could be associated with the amount of centromeric DNA sequence^20–23^. Other independent reports revealed a statistically significant higher number of small chr21 α-satellite HOR arrays in individuals with the syndrome compared to controls^27^, suggested an association of small chr21 α-satellite HOR arrays and MMIEs^42^, and proposed that individuals with T21 generally harbor small chr21 centromeres^27^. Nevertheless, a 2003 study explored the relationship between the size of α-satellite HOR arrays on chr21 centromeres and repressed recombination, one of the known phenomena that correlate with nondisjunction, and concluded that the features are independent^43^.

In our investigation, chr21 centromeres showed remarkable variability in length, α-satellite HOR array structure, and CDR patterns within the human population (Figure 4), suggesting that chr21 is amongst the top four most variable centromeres in the human genome^7^. We show that small centromeres are found at low frequency in individuals with T21 (3/24; 3 centromeres smaller than 500 kbp in 3 individuals, 2 of which of maternal origin) and that T21 centromere size distribution is comparable to that of the general population (Figure 5a). Despite being present at low frequency in the general population (10%), our data do not support the observation that small centromeres alone constitute a risk factor for T21 nondisjunction. Rather, our results suggest an alternative hypothesis, where centromere size asymmetry may be a more important consideration and possibly one of the features that contribute to suppressed recombination and nondisjunction.

Small centromeres might have been previously reported in children with T21, as they make size asymmetry more likely than larger centromeres. Our results support other observations^44^, which noted that small centromeres are not exclusive of individuals with T21; contrarily, extreme homologous chr21 centromere size asymmetry (>10-fold) appears to be absent in individuals without the syndrome (Figure 5b**; Supplementary** Figure 10). We report two cases in families that include an individual with T21 with similar features, but none in the general population.

Moreover, our phylogenetic analysis (Figure 4a) shows that rapid changes of chr21 centromere size are possible and might facilitate the occurrence of centromere size asymmetry in the population. We speculate that an extreme difference between chr21 centromere sizes makes homologous pairing less efficient for example by complicating meiotic synapsis across the entire centromere length. There is evidence demonstrating that larger domains of homology are essential for synapsis formation and/or recombination in complex organisms such as *C.elegans*, *Drosophila*, and mice but not in simpler organisms like yeast^45–49^. Nevertheless, more experimental data are needed to test how asymmetry impacts meiosis, recombination, and chromosome segregation.

The families (AG167 and 7210079) showing extreme centromere size asymmetry (10.7- and 19.4-fold) have other features in common, suggesting a common T21 aetiology. For instance, they carry the smallest chr21 centromeres ever characterized in females (143 and 181 kbp), both exhibit a transgenerational CDR change on the longer homologous centromere, and both mothers had a similar age at time of childbirth (29 and 27 years old). Another family (7210223) exhibited a transgenerational CDR change. In this case, we do not observe an extreme centromere size asymmetry (2.6-fold), and the methylation change occurred on the smaller haplotype (587 kbp). Mother 7210223.20 was 22 at childbirth and we did not observe any other unusual centromeric features, thus suggesting an alternative mechanism underlying T21.

In this study we also investigated the amount of centromeric proteins forming the kinetochore complex on each of the chr21 centromeres of family AG167. Our IF-FISH data (Supplementary Figure 3) suggest that none of the chr21 centromeres have a significantly lower amount of CENP-A or CENP-C proteins, arguing against the hypothesis that MMIEs are caused by malfunctions in the kinetochore complex. It is unclear how the kinetochore function is impacted by the altered CDRs observed in three of the children with T21 and we cannot determine whether the poorly defined CDRs were already present in the maternal germ cells or developed later in the children after establishment of T21.

The first family with T21 (AG167) centromere showed epigenetic transgenerational differences between maternal and child centromeres in the absence of genetic changes. Specifically, we observed significant differences in CDR size of child’s maternal H2 and paternal H3 CDRs as well as a positional shift of ∼11 kbp and 25 kbp towards the p-arm in the child for the former and the latter, respectively (Figure 2c), despite the fact that the sequence and structure between parent and child were practically identical (Supplementary Figure 5). This reinforces the notion that CDRs and kinetochore position are not defined by the primary sequence but can change location potentially even within a single generation. This is also supported by two population haplotypes, HG03710-H1 and HG02647-H2, which both carry their CDRs closer to the p-arm yet differ radically in their HOR array monomers with 5-monomers and 4-monomers corresponding to older and younger repeats, respectively.

The second cohort of families with T21 also allowed us to compare CDR differences between cell lines and a primary tissue like blood. Little is known regarding the stability of CDRs somatically; however, our limited data on multiple samples demonstrate that CDRs position between blood and cell lines remain positionally conserved. The only detected difference is a generally higher level of methylation in the blood. This could be due to multiple reasons. First, comparison of methylation callers (Supplementary Figure 8) on cell line data indicates that PacBio basecallers (Jasmine and especially the older Primrose) detect a slightly higher level of methylation. Also, HiFi reads might be mapping less precisely than the much longer UL-ONT reads. Lastly, it is possible that some highly specialized cells in the blood, which are generated from the progenitors in the bone marrow and do not typically undergo mitosis^50^, might lose epigenetic signatures over time like those associated with CDRs.

There are several limitations in this study. First and foremost, the observation of asymmetry rests on relatively few observations in part because the sequence and assembly of centromeres is still a costly and time-consuming endeavor. Moreover, we focused here on individuals with T21 conceived by younger mothers (age range: 20-34) despite the known, strong association of nondisjunction with advanced maternal age. It will be important to pull apart these confounders by assessing centromere asymmetry in even younger and older mothers and incorporating the maternal age effect. Larger T21 population-based studies are, therefore, required to confirm this finding as well as evaluating families with MMIEs and MMIIEs. Large cohorts with short-read sequence data, such as the Atlanta or the National Down Syndrome Projects^36,37^, have been assembled and imputation of chr21 centromere length in these cohorts once a large enough pangenome has been amassed may be possible instead of direct sequencing. Finally, future studies should evaluate not only centromere sequence and methylation using long-read sequencing data but also evaluate the composition and location of the kinetochore complex as it relates to epigenetic differences of the CDR.

## Acknowledgments

This work was supported, in part, by US National Institutes of Health (NIH) grants R01HG010169 and U24HG007497 (to E.E.E.) and R00GM147352 (to G.A.L.). E.E.E. is an investigator of the Howard Hughes Medical Institute. This article is subject to HHMI’s Open Access to Publications policy. HHMI lab heads have previously granted a nonexclusive CC BY 4.0 license to the public and a sublicensable license to HHMI in their research articles. Pursuant to those licenses, the author-accepted manuscript of this article can be made freely available under a CC BY 4.0 license immediately upon publication.

## MATERIAL AND METHODS

### Sample recruitment and data collection

The following lymphoblastoid cell lines were obtained from the NIGMS Human Genetic Cell Repository at the Coriell Institute for Medical Research: proband (AG16777), mother (AG16778), and father (AG16782). The proband is described as a mosaic with karyotype: 47,XX,+21[21]/47,XX,+21,t(21;22)(q22;q13)[29] with 10% of the cells examined showing random chromosome loss and the karyotype confirming the diagnosis. Parents do not have Down syndrome and have a normal karyotype. The proband was 43 years old at sampling, while mother and father were 72 and 74, respectively. We obtained lymphoblastoid cell lines from Emory University biobank for seven probands with MMIE T21 and six mothers. Blood samples were also obtained for the same probands.

### PacBio HiFi sequencing

PacBio HiFi data were generated per manufacturer’s recommendations. Briefly, high-molecular-weight (HMW) DNA was extracted from frozen blood or cultured lymphoblasts using the Monarch® HMW DNA Extraction Kit for Cells & Blood (New England Biolabs, T3050L). At all steps, quantification was performed with Qubit dsDNA HS (Thermo Fisher, Q32854) measured on DS-11 FX (Denovix) and size distribution checked using FEMTO Pulse (Agilent, M5330AA & FP-1002-0275.) HMW DNA was sheared with Megaruptor 3 (Diagenode, B06010003 & E07010003) using settings 28/31 or 28/30 (depending on original length distribution) and used to generate PacBio HiFi libraries via the SMRTbell Prep Kit 3.0 (PacBio, 102-182-700). Size selection was performed with Pippin HT using a high-pass cutoff of 17 kbp (Sage Science, HTP0001 & HPE7510.) Samples AG16778, AG16782, and NG16777 were sequenced on the Sequel II platform on SMRT Cells 8M (PacBio, 101-389-001) using Sequel II Sequencing Chemistry 3.2 (PacBio,102-333-300) with 2-hour pre-extension and 30-hour movies, aiming for a minimum estimated coverage of 30× in PacBio HiFi reads for parents (3 SMRT Cells per sample) and 60× for the proband (5 SMRT Cells, assuming a genome size of 3.1 Gbp). The remaining samples were sequenced on the Revio platform on SMRT Cells 25M (PacBio, 102-817-900) with diffusion loading and 24-hour movies or Adaptive Loading and 30-hour movies, aiming for a minimum estimated coverage of 30× in PacBio HiFi reads for the parents (1-1.5 SMRT Cells per sample) and 60× for the probands (2-2.5 SMRT Cells, assuming a genome size of 3.1 Gbp).

### UL-ONT sequencing

Ultra-high molecular weight gDNA was extracted from the lymphoblastoid cell lines according to a previously published protocol. Briefly, 3-5 × 10^7 cells were lysed in a buffer containing 10 mM Tris-Cl (pH 8.0), 0.1 M EDTA (pH 8.0), 0.5% w/v SDS, and 20 mg/mL Rnase A for 1 hour at 37°C. 200 ug/mL Proteinase K was added, and the solution was incubated at 50°C for 2 hours. DNA was purified via two rounds of 25:24:1 phenol-chloroform-isoamyl alcohol extraction followed by ethanol precipitation. Precipitated DNA was solubilized in 10 mM Tris (pH 8.0) containing 0.02% Triton X-100 at 4°C for two days. Libraries were constructed using the Ultra-Long DNA Sequencing Kit (ONT, SQK-ULK001 or SQK-ULK114) following the manufacturer’s protocol. 75 uL of library was loaded onto a primed FLO-PRO002 R9.4.1 or FLO-PRO114M R10.4.1 flow cells for sequencing on the PromethION, with two nuclease washes and reloads after 24 and 48 hours of sequencing. Sequence reads >100 kbp in length were classified as ultra-long (UL). All ONT data were base called using guppy (v6.3.7 or newer) or dorado (v0.4.2 or newer) with the SUP model.

### Genome assembly

Proband AG16777 genome assembly was generated with the Verkko^32^ (v1.2) (https://github.com/marbl/verkko) workflow using HiFi sequencing data at ∼59× and UL-NT (>100 kbp) data at ∼23× coverage following the developer instructions. Parental assemblies (AG16778 and AG16782) were generated with both Verkko^32^ (v1.4) (https://github.com/marbl/verkko) and hifiasm^51^ (v0.19.5) (https://github.com/chhylp123/hifiasm) using HiFi data at ∼37× for the mother and 40× for the father and UL-ONT data at 25× and 12×, respectively. HiFi developer instructions were followed. Similarly, genome assemblies for the second cohort of probands with T21 and their mothers were generated using Verkko^32^ (2.0 or newer) and hifiasm^51^ (v0.19.8 or newer).

### Cell culture, slide preparation and Immunofluorescence (IF) combined with fluorescence *in situ* hybridization (FISH)

AG16777, AG16778 and AG16782 cells were cultured in RPMI-1640 medium supplemented with 16% fetal bovine serum, 1% L-glutamine, and 1% penicillin– streptomycin. Metaphase and interphase nuclei undergoing immunofluorescence with mouse CENP-A monoclonal antibody (Enzo, ADI-KAM-CC006-E) were prepared by blocking the cells in metaphase with colcemid, followed by washing in phosphate-buffered saline (1× PBS). The cells were then counted and resuspended in a hypotonic buffer (75 mM KCl: 0.8% NaCitrate: 3 mM CaCl2: 1.5 mM MgCl2, 1:1:1 ratio) for 15 minutes to reach a final concentration of 5 x 10⁵ cells/mL. A 0.5 mL aliquot of the cell suspension was then cytocentrifuged onto Superfrost Plus slides at 1,500 rpm for 5 minutes using a Shandon Cytospin 3 (Thermo Scientific). The slides were immediately used for immunofluorescence. For slides undergoing immunofluorescence with mouse CENP-C monoclonal antibody (Abcam, ab50974), colcemid-treated cells were incubated in hypotonic solution (0,56% KCl) for 30 min and prefixed by adding methanol:acetic acid (3:1) fixative solution. Prefixed cells were collected by centrifugation and then fixed in methanol:acetic acid (3:1). Spreads were dropped on a glass slide and incubated at RT overnight.

Immunofluorescence was performed on the metaphase spreads using mouse CENP-C and CENP-A monoclonal antibodies as previously described with minor modifications^52^. In brief, each slide was rehydrated by immersion in 1× PBS-azide (10 mM NaPO4, pH 7.4, 0.15 M NaCl, 1 mM EGTA and 0.01% NaN3) for 15 min at room temperature. Chromosomes were then swollen by washing the slides (three times, 2 min each) with 1× TEEN (1 mM triethanolamine-HCl, pH 8.5, 0.2 mM NaEDTA, and 25 mM NaCl), 0.5% Triton X-100 and 0.1% BSA. The primary antibodies were diluted to a final concentration of 0,001 μg/μl in the same solution and then added (100 μl) onto the slides. Each slide was incubated for 2 h at 37 °C. Excess of primary antibody was removed by washing the slides at room temperature (three times, 2, 5 and 3 min each) with 1× KB buffer (10 mM Tris-HCl, pH 7.7, 0.15 M NaCl and 0.1% BSA). A goat anti-mouse IgG secondary antibody conjugated to FITC (Abcam, ab6785) was diluted 1:100 in the same solution, and 100 μl was then added to the slides that were then incubated for 45 min at 37 °C in a dark chamber. After incubation with the secondary antibody, the slides were washed once with 1× KB for 2 min, prefixed with 4% paraformaldehyde in 1× KB for 45 min at room temperature, washed with dH2O by immersion for 10 min at room temperature, and fixed with methanol and acetic acid (3:1) for 15 min. FISH was then performed using a plasmid containing the α-satellite sequence shared between chromosomes 13 and 21 (pZ21A) directly labelled by nick-translation with Cy3-dUTP (Enzo, 42501) according to a standard procedure with minor modifications^53^. In brief, 100 ng of labelled probe was used for the FISH experiments; DNA denaturation was performed at 70 °C for 8 min and hybridization at 37 °C in 2× SSC, 50% (v/v) formamide, 10% (w/v) dextran sulphate and 3 mg sonicated salmon sperm DNA, in a volume of 10 μl. Post-hybridization washing was performed under high stringency conditions: at 60 °C in 0.1× SSC (three times, 5 min each). Nuclei and chromosome metaphases were simultaneously DAPI-stained. Digital images were obtained using a Leica DMRXA2 epifluorescence microscope equipped with a cooled CCD camera (Princeton Instruments). DAPI, Cy3 and fluorescein fluorescence signals, detected with specific filters, were recorded separately as grayscale images. Pseudocolouring and merging of images were performed using ImageJ (v.1.53k). The CENP-C fluorescence intensity at chr21 centromeres was measured on 13 metaphases for each sample using a modified version of the ImageJ macro, CRaQ^54^. Signals from chr21 were isolated, and a 7 × 7 pixel box was placed around the centroid position of each signal. The pixel intensity within each box was measured and normalized dividing it by the lower chr21 signal intensity for each analyzed metaphase. Plots were then obtained using the ggplot2^55^ package in R^56^.

### Centromere characterization and validation

Assemblies of individuals with T21 and from the general population were aligned against the T2T-CHM13^8^ (v2.0) reference genome with minimap2^57^ (v2.24) (https://github.com/lh3/minimap2) using the following options “-t {threads} -I 15G -a –eqx -x asm20 -s 5000”. Fasta sequences from the contigs mapping against chr21 centromere (chr21:10700000-11650000) and flanking regions were extracted. RepeatMasker^58^ (v4.1.0) was used to identify the regions containing the α-satellite array (marked as “ALR/Alpha’’) and HumAS-HMMER (https://github.com/fedorrik/HumAS-HMMER_for_AnVIL) to determine the α-satellite HOR array composition with the following command: hmmer-run.sh {path_to_directory_with_fasta} AS-HORs-hmmer3.0-170921.hmm {threads}. For the contigs aligning to the reference genome in the minus orientation, we converted the fasta sequences to their reverse complement before running the analysis. The outputs from HumAS-HMMER analyses were visualized with R^56^ (v4.2.2) using the ggplot2^55^ package.

Another 44 population chr21 centromere sizes were obtained from the recently published HGSVC data^35^. Additionally, 115 female and 101 male samples from the HPRC year 2 release were processed and validated with CenMap (v0.3.1) (https://github.com/logsdon-lab/CenMAP), a newly developed pipeline for centromere mapping and annotation, to calculate their chr21 centromere sizes.

We validated each centromeric contig sequence aligning each sample’s HiFi reads to their own genome assembly using pbmm2 (v1.1.0) (https://github.com/PacificBiosciences/pbmm2) and showing uniform read coverage across the centromeric region with NucFreq^59^ (https://github.com/mrvollger/NucFreq). To confirm that the extracted centromeric contig sequences represented chr21 centromere repeats, we compared them to the α-satellite monomer units extracted from the T2T-CHM13^8^ reference genome using StringDecomposer^60^ (https://github.com/ablab/stringdecomposer) and showed they have high identity (>90%).

Finally, we generated pairwise sequence identity heatmaps of each centromere using StainedGlass as previously described^33^ (https://github.com/mrvollger/StainedGlass).

### Supplementary note

We were not able to characterize both chr21 centromeres in 35 females and 30 males from the HPRC year 2 release. In these samples, one or none of the chr21 centromeres was assembled, or the haplotypes could not be reliably assigned to chr21 or chr13, which typically has much larger centromeres. This is common when the genomic contigs containing the centromeric sequences are not sufficiently extended to encompass the pericentromeric regions of both p- and q-arms and highlights the difficulty of sequencing and assembling large centromeres. As an example, two female samples (HG00320 and HG04184) carried a small chr21 centromere (228 and 263 kbp), but their putative homologs were either incomplete or contained errors, with ambiguous assignments to chr21 or chr13. Some of these contigs contained large α-satellite arrays, but despite further analysis, we were unable to conclusively determine whether these putative homologs represented chr21, chr13 or a mix of the two centromeres. Consequently, they were excluded. It is worth noting though that even with two cases of extreme size asymmetry in the female population cohort, we would still have a significant difference in the number of cases observed in the two groups (population samples = 80; Fisher’s exact test p-value = 0.031).

### Centromere alignments and dotplots

AG16777 proband and parental centromere sequences were aligned using minimap2^57^ (v2.26) and the following command: minimap2 -cx asm5 {proband_centromeric_fasta} {corresponding_parental_centromeric_fasta} > alignment.paf. The results in .paf format were visualized with R^56^ (v4.2.2) using the ggplot2^55^ package. Proband H3 and father H3 AluY centromeric insertions with 10 kbp padding sequence on both sides were aligned with BLAST^61^.

### CpG methylation and CDR analysis

All ONT sample data were basecalled with guppy v6.3.7 or newer to generate BAM files containing methylation tags linked to each read. HiFi data were base called using CCS (v.6.4.0 or newer) (https://github.com/PacificBiosciences/ccs), Primrose (v1.4.0 or newer) or (Jasmine (v2.0.0 or newer) (https://github.com/PacificBiosciences/jasmine), which generated the equivalent BAM files containing the methylation tags linked to each read. Jasmine 5mC predictions were used as a validation for the centromeric methylation profiles. Briefly, we extracted the reads for each sample and aligned them to their respective genome assembly using Winnowmap^62^ (v2.03 or newer). We then used modbam2bed (https://github.com/epi2me-labs/modbam2bed) to obtain bedMethyl files.

Using the bedMethyl files as an input and an early developer version of CDR-Finder^17^, we identified the CDRs for each centromere. The tool first divides the region of interest (the centromeric α-satellite arrays with padding sequence on both sides, e.g., coordinates for Proband H1: unassigned-0003018:0-2000000) in sequential 5 kbp bins and calculates their average methylation frequency (if a bin does not have an assigned value, the bin is excluded). It then computes the median CpG methylation frequency for each window containing α-satellite sequences (based on RepeatMasker^58^ output). It selects the bins with a lower methylation frequency than the median of the region and merges consecutive bins into candidate CDRs.

These candidate CDRs are then evaluated on two criteria: whether or not they have flanking windows on either side with greater than the maximum percent methylation (defined as within 1 standard deviation of the maximum) for the entire region and if they clear a user-specified size threshold (25 kbp). Each call is annotated with a confidence label: “high confidence” if it clears both criteria or “low confidence” if it does not. The low-confidence labels are appended with either “neighbor_peaks” for the first criterion and/or “size” for the second criterion. In small centromeres where the median can be quite low due to the CDR taking up the majority of the centromeric sequence space, the initial search algorithm of taking windows which are below the median is flipped to identify the windows that are above the median and thus not part of the CDR. After removing these windows from consideration, what remains are candidate CDRs, and the algorithm continues as normal. All CDR calls were manually confirmed inspecting the read alignments. False positives at the boundaries of the centromeres as well as calls not confirmed by the reads and due to low coverage were excluded.

For each AG167 family CDR, we calculated the percentage of methylated CpGs (defined as having ≥25% methylated CpGs at the same position across all UL-ONT reads) within the total CDR window (from start of the first CDR window to the end of the last CDR window). Formula = (number of methylated CpGs within the CDR window) * 100 / (number of CpGs within the CDR window).

### Contig fixing and patching

Centromeric contigs that showed misassemblies or collapses during NucFreq^59^ analysis (above) were corrected following this procedure. Sequencing reads mapping to the misassembly/collapse and flanking regions were extracted and locally assembled using the module Canu from the tool HiCanu^63^. The local assemblies were validated using NucFreq^59^ as described previously and inserted in the original assembly.

Verkko^32^ genome assemblies containing errors were corrected with an alternative assembly generated via hifiasm^51^ or vice versa. Briefly, we generated assemblies of each genome with hifiasm^51^ (v0.19.5 or newer) and the following command: “hifiasm -o {output} -t {threads} –ul {ultra-long read fastq} {hifi reads fastq}”. Then, we aligned the hifiasm assemblies to the Verkko ones with minimap2^57^ (v2.24) and the following command: minimap2 -t {threads} -I 15G -a –eqx -x asm20 -s 5000. We identified the corresponding locations of the centromeric regions in each assembly and then determined the α-satellite HOR array composition of each with HumAS-HMMER (https://github.com/fedorrik/HumAS-HMMER_for_AnVIL) and the following command: hmmer-run.sh {path_to_directory_with_fasta} AS-HORs-hmmer3.0-170921.hmm {threads}. We compared the α-satellite HOR compositions of each array and then replaced erroneous sequences in the Verkko assembly with the corresponding sequence in the hifiasm assembly (or vice versa). To ensure that we had resolved the assembly error, we aligned PacBio HiFi reads from the same genome to the new assembly and then ran NucFreq^59^ to validate the assembly.

### Phylogenetic analysis

For the phylogenetic reconstruction of chr21 centromere haplotypes, we first aligned a chimpanzee assembly to the T2T-CHM13^8^ reference genome using Minimap2^57^ with the following options “-t {threads} -I 15G -x asm5 -a –secondary=no”. We identified a chr21 pericentromeric region on the q-arm where only two chimpanzee contigs mapped (chr21:11855000-11875000). The chimpanzee alignment was used as an outgroup for the analysis. We aligned all the human assemblies available against the T2T-CHM13^8^ reference genome as described above and selected only the samples with chr21 centromeric contigs spanning the selected pericentromeric window. We then extracted the sequences mapping to the pericentromeric region of interest of each sample. Using these sequences, we generated a multi-alignment with the tool MAFFT^64,65^ (v7.453) using these options: “–auto –thread {threads}”. The maximum-likelihood phylogeny tree was constructed with IQ-TREE^66^ (v2.1.2) with the following options: “-T {threads} -m MFB -B 1000”. Divergence time amongst haplotypes were estimated with IQ-TREE^66^ (options: “–keep-ident –date-tip 0 –date-ci 100 -redo”) using the MAFFT multi-alignment, the consensus tree, and a time estimate file containing the known divergence time between human and chimpanzee (6.4 million years) as inputs. The resulting tree was visualized with iTOL^67^ and FigTree (https://github.com/rambaut/figtree).

## Data availability

Sequencing data were submitted to the database of Genotypes and Phenotypes (dbGaP) for general research use and public access. The phs accession ID is: phs003761.v1.p1.

## Supplementary Figures

**Supplementary Figure 1.**
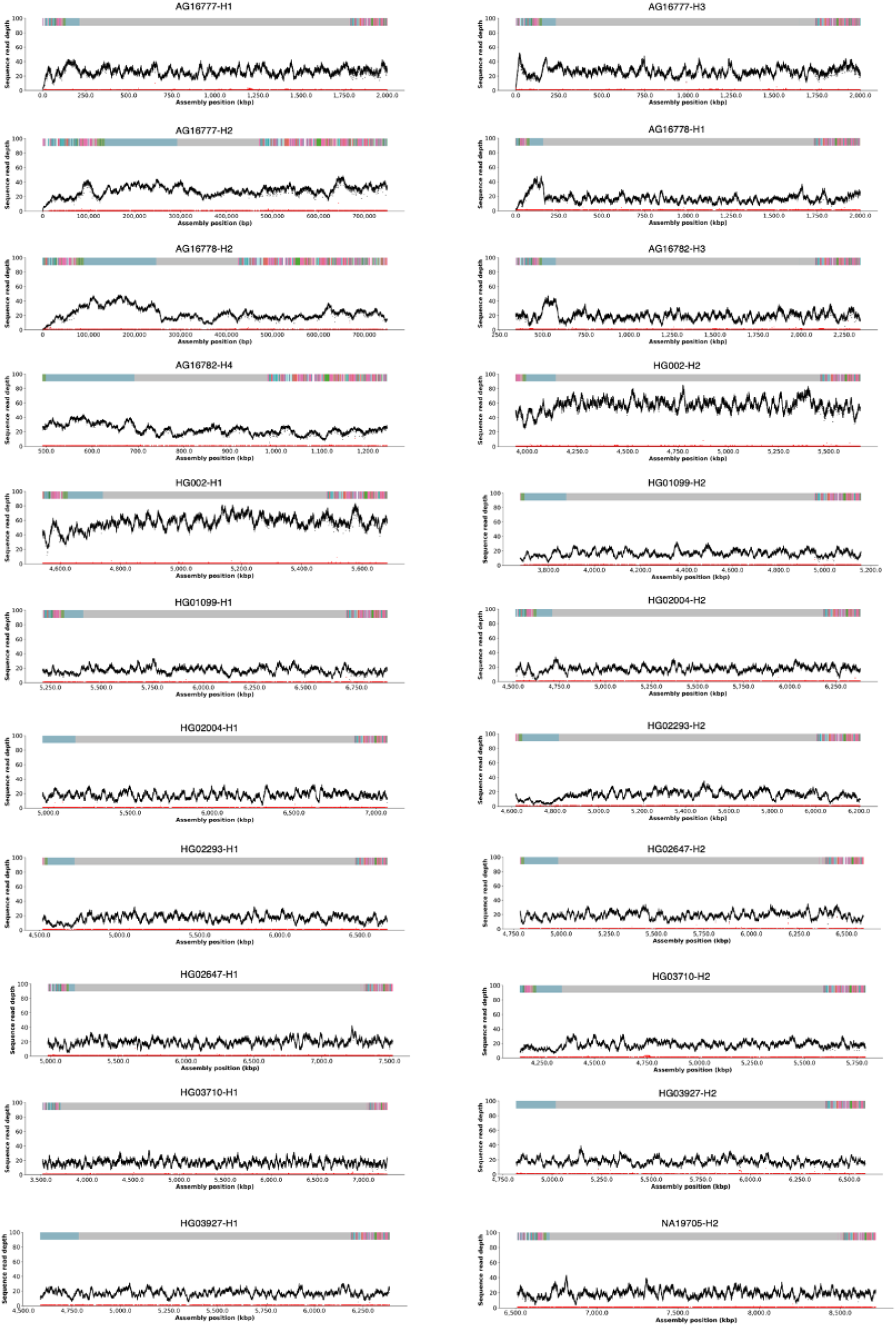
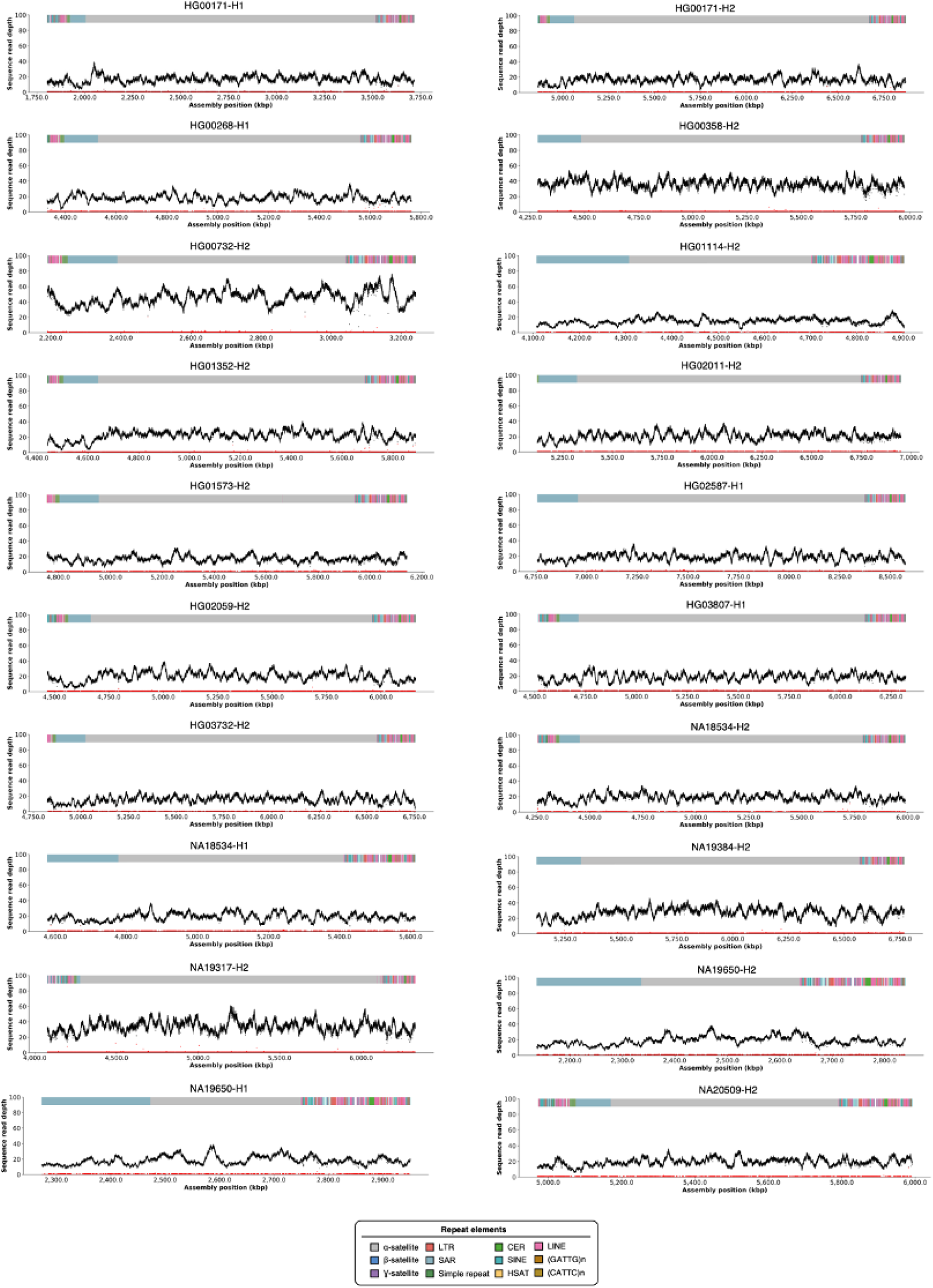
NucFreq validation of all the chromosome 21 (chr21) centromeres described in the family AG167 and population samples.

**Supplementary Figure 2.**
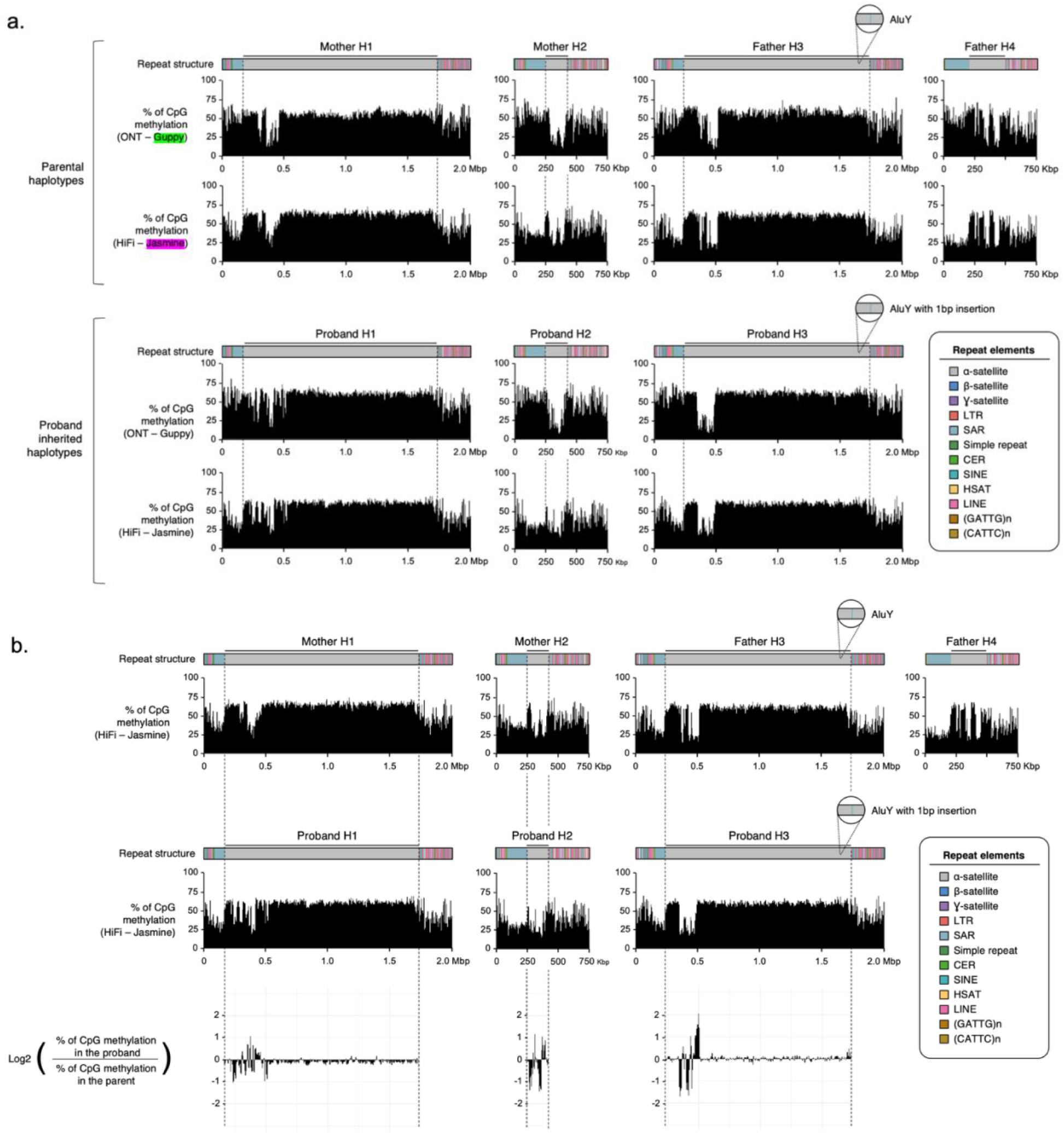
**a)** Comparison of the ONT- and HiFi-based methylation profiles in all family AG167 chr21 centromeres. **b)** Comparison between proband and parental chr21 centromere methylation profiles based on the HiFi data.

**Supplementary Figure 3.**
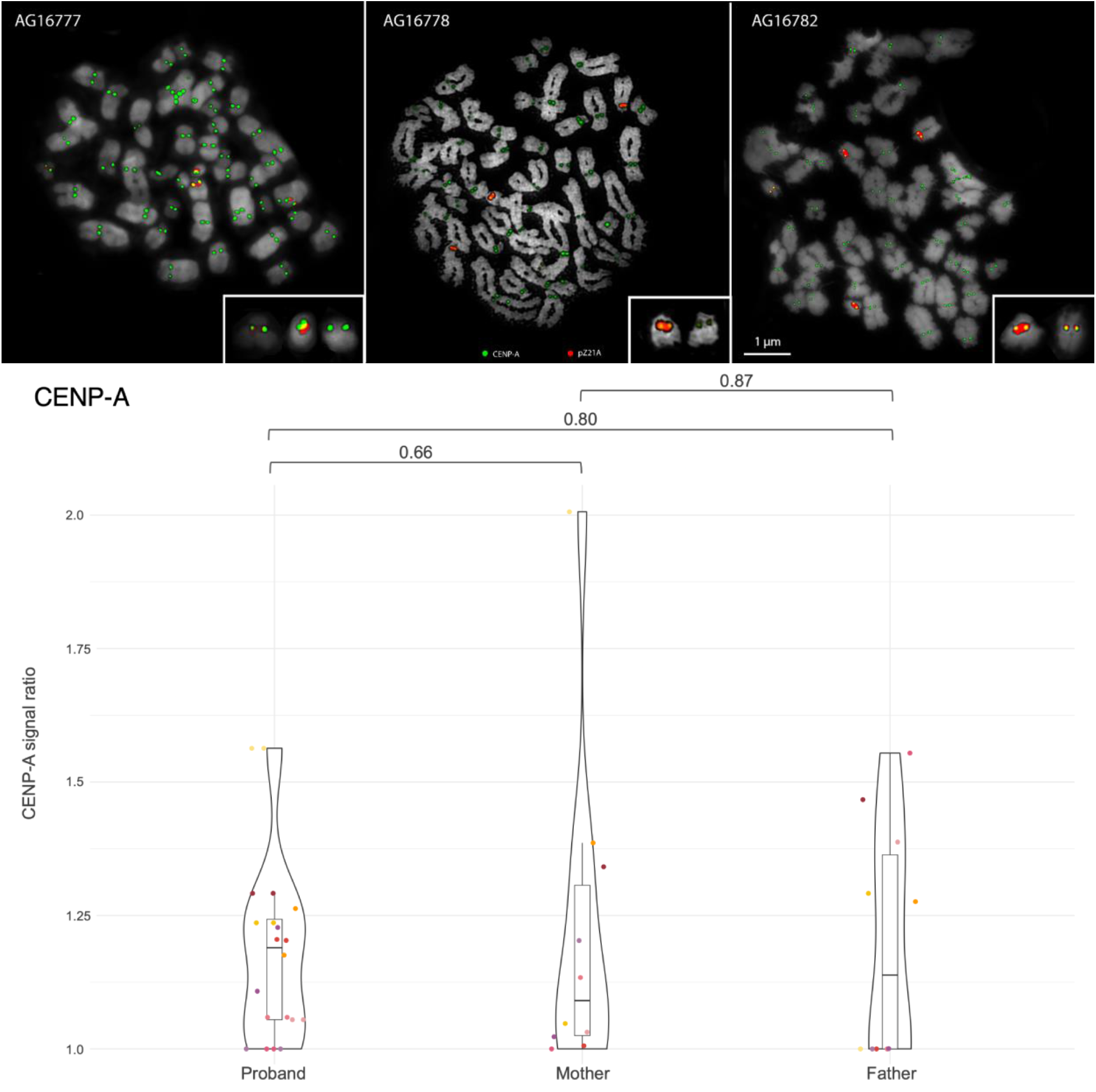
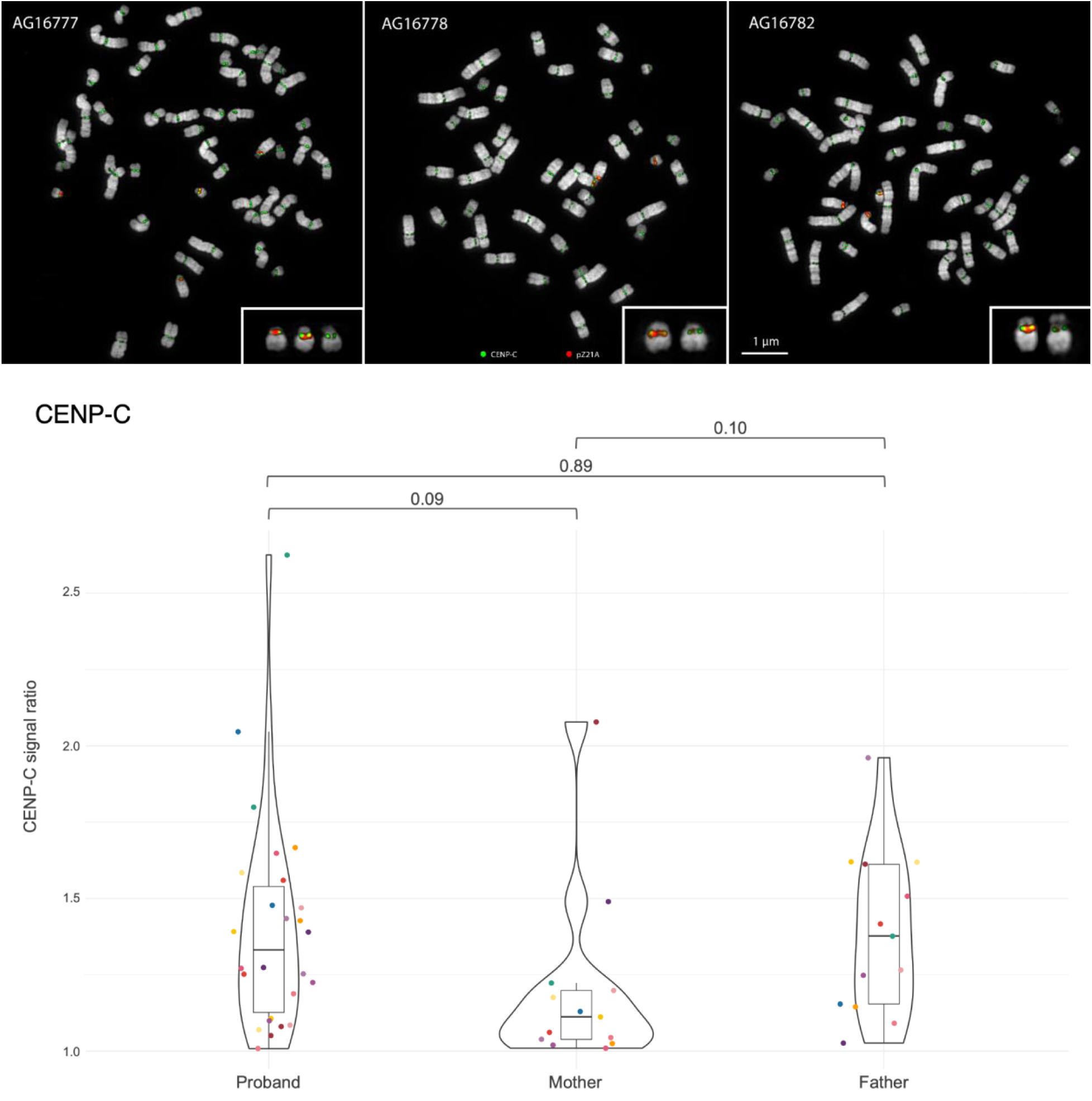
CENP-A and CENP-C detection and quantification in proband AG16777, mother AG16778, and father AG16782. IF-FISH on metaphase chromosome spreads from all members of the AG167 family, with fluorescent probes targeting the chr13/21 α-satellite array (red) and CENP-A/CENP-C (green). All chr21 are highlighted in the boxes. A weaker α-satellite array probe signal on one of the chr21 centromeres of proband and mother is due to the smaller centromeric α-satellite HOR array present in each of these individuals. The violin plots represent the CENP-A and CENP-C signal ratios for each member of the family. Each data point represents a metaphase from which the values of CENP-A/CENP-C protein signals were quantified. Each signal value was divided by the homolog lowest value to obtain the ratio and self-comparisons were removed. Signal ratio of 1 indicates an identical value between homolog signals. T-test p-values are indicated at the top for each comparison.

**Supplementary Figure 4.**
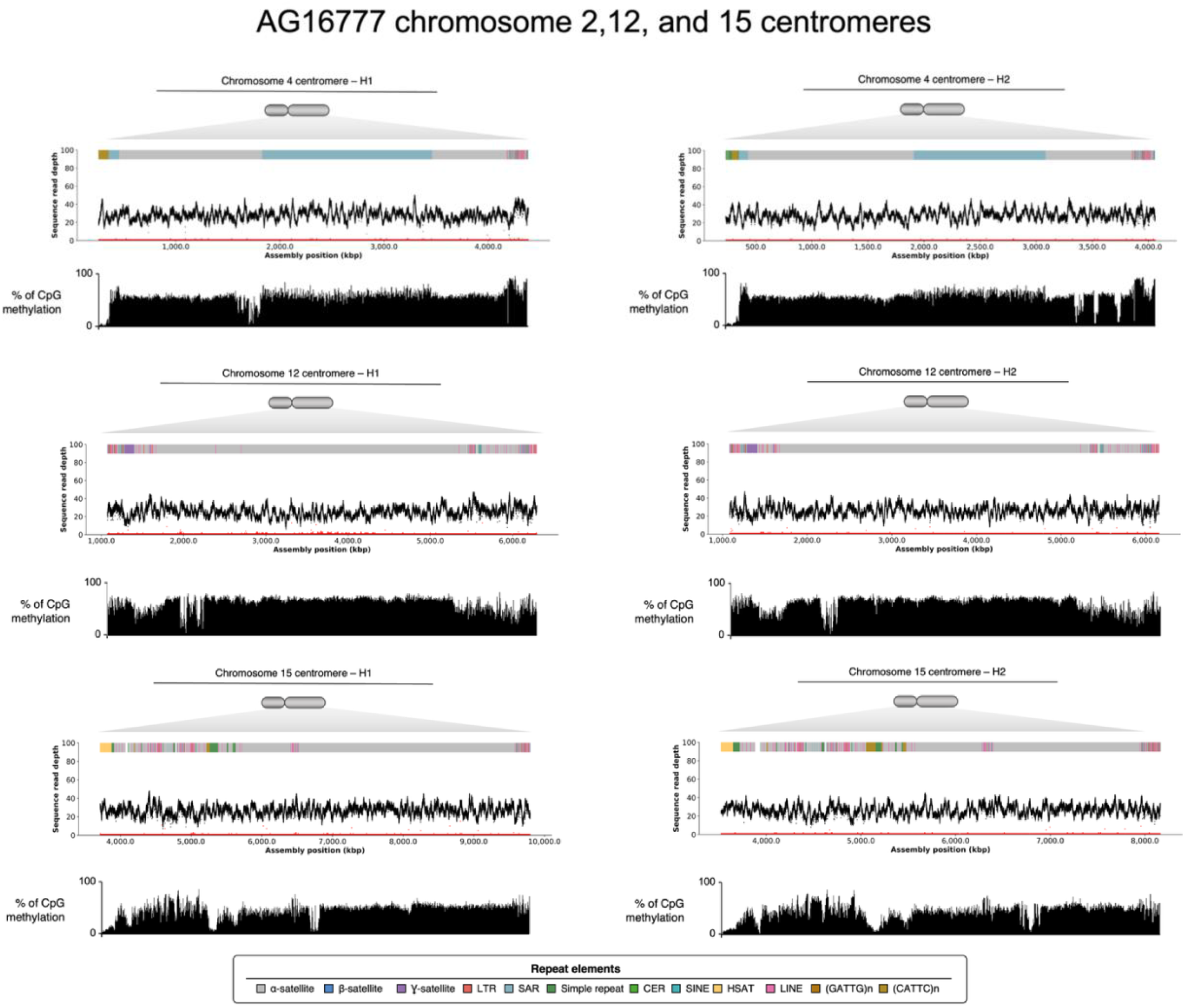
Repeat structure, NucFreq plot validation, methylation profile of chromosome 4, 12, and 15 centromeres in proband AG16777.

**Supplementary Figure 5.**
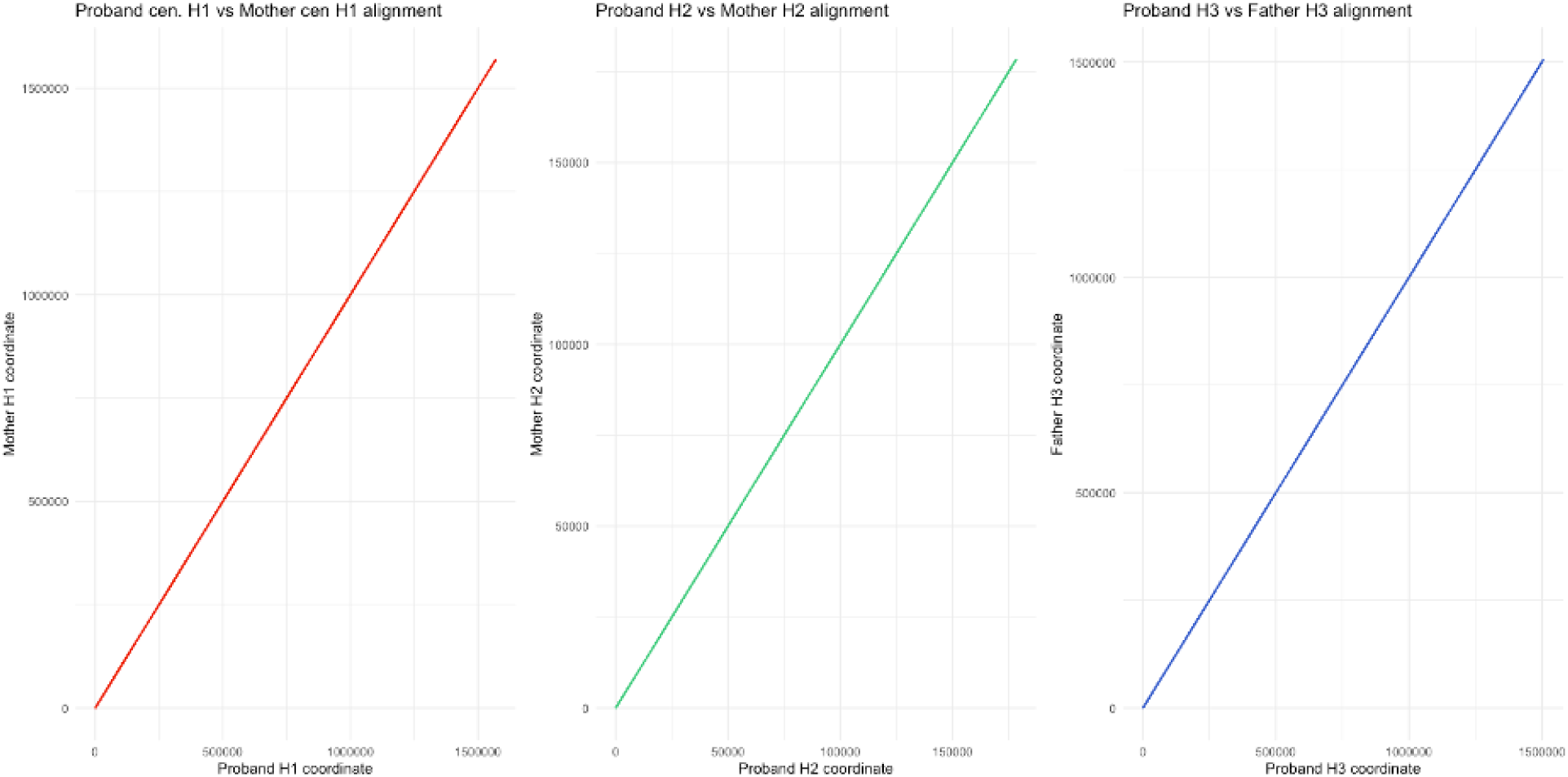
Dotplots comparing AG167 proband and parental centromeric sequences haplotypes. The plots represent the alignments of proband and parental centromeric α-satellite array sequences.

**Supplementary Figure 6.**
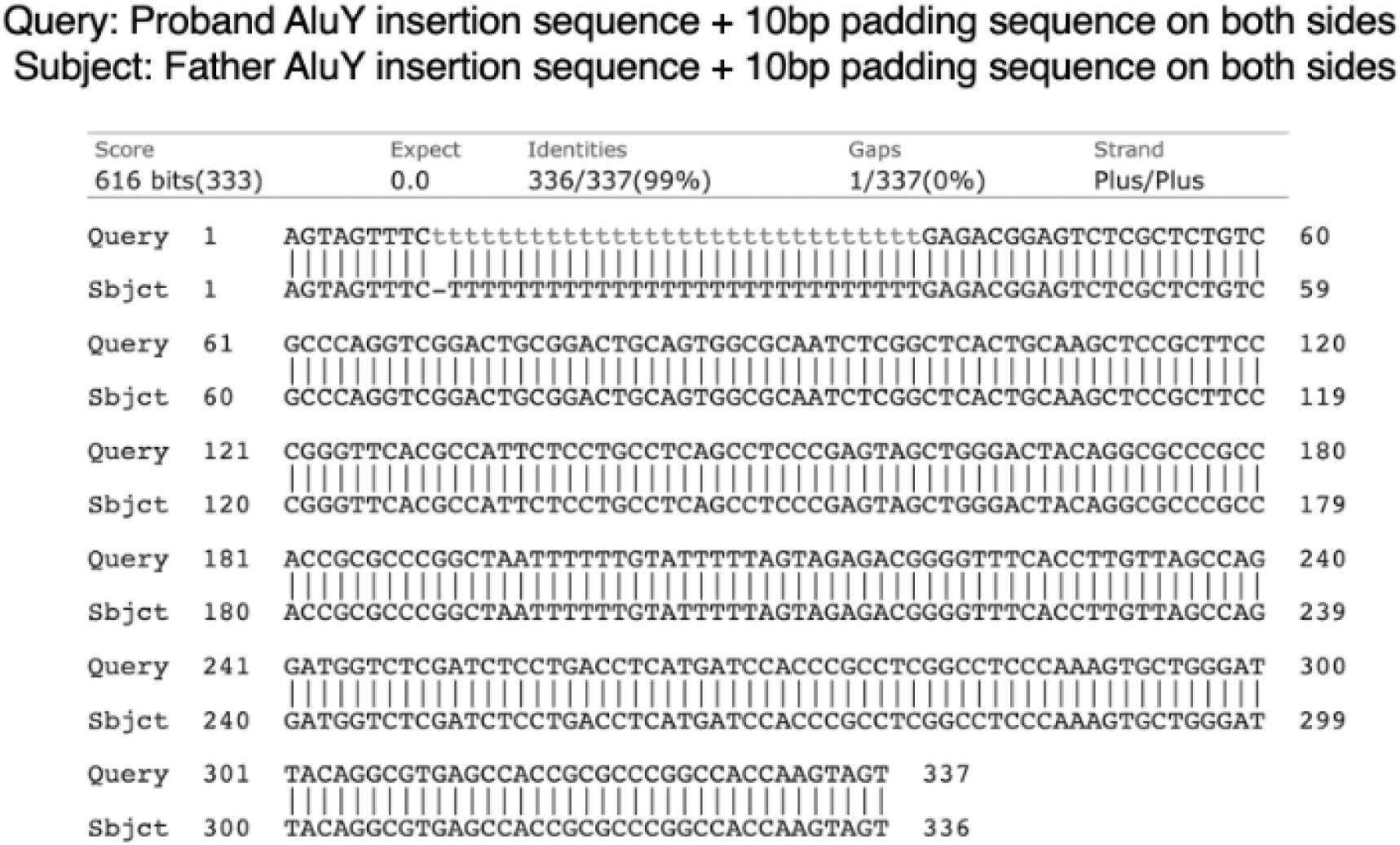
BLAST-based pairwise alignment of the AluY sequences inserted in AG167 proband H3 and father H3 centromere sequences.

**Supplementary Figure 7.**
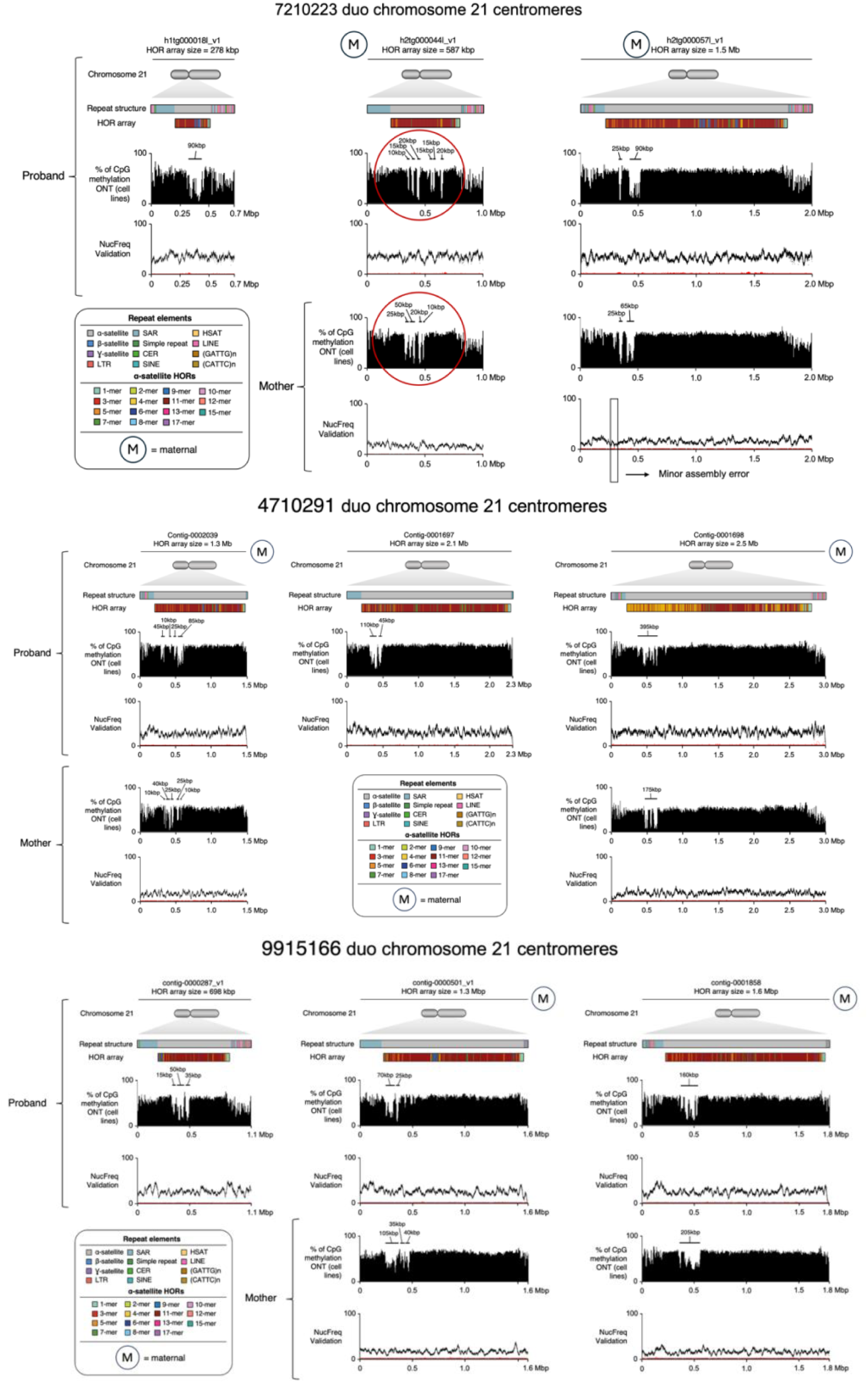
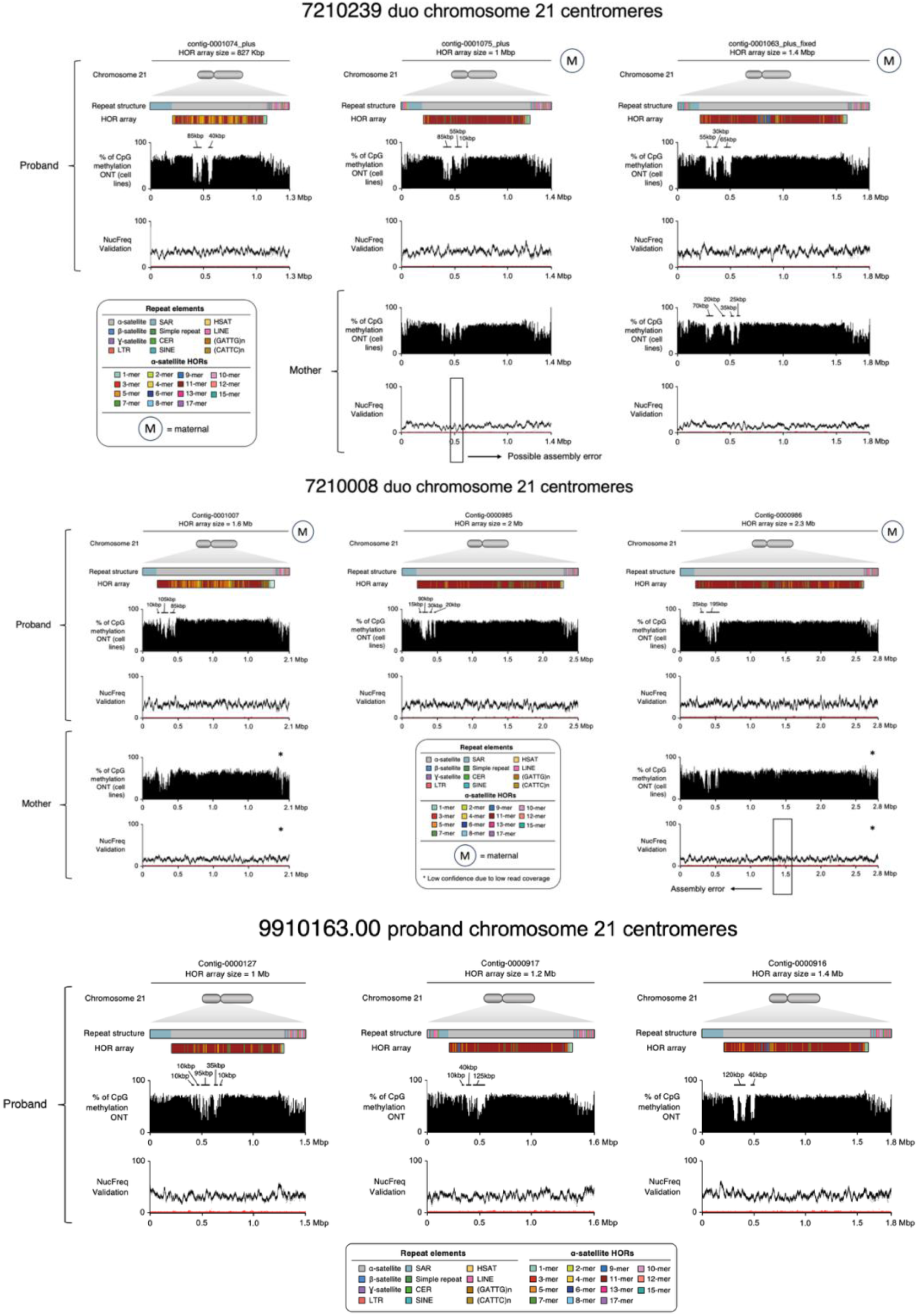
Centromeres of individuals with T21 from families that do not exhibit extreme maternal homolog size asymmetry. Repeat structures, HORs and methylation profiles are shown. 7210223 duo is the only one showing transgenerational methylation changes.

**Supplementary Figure 8.**
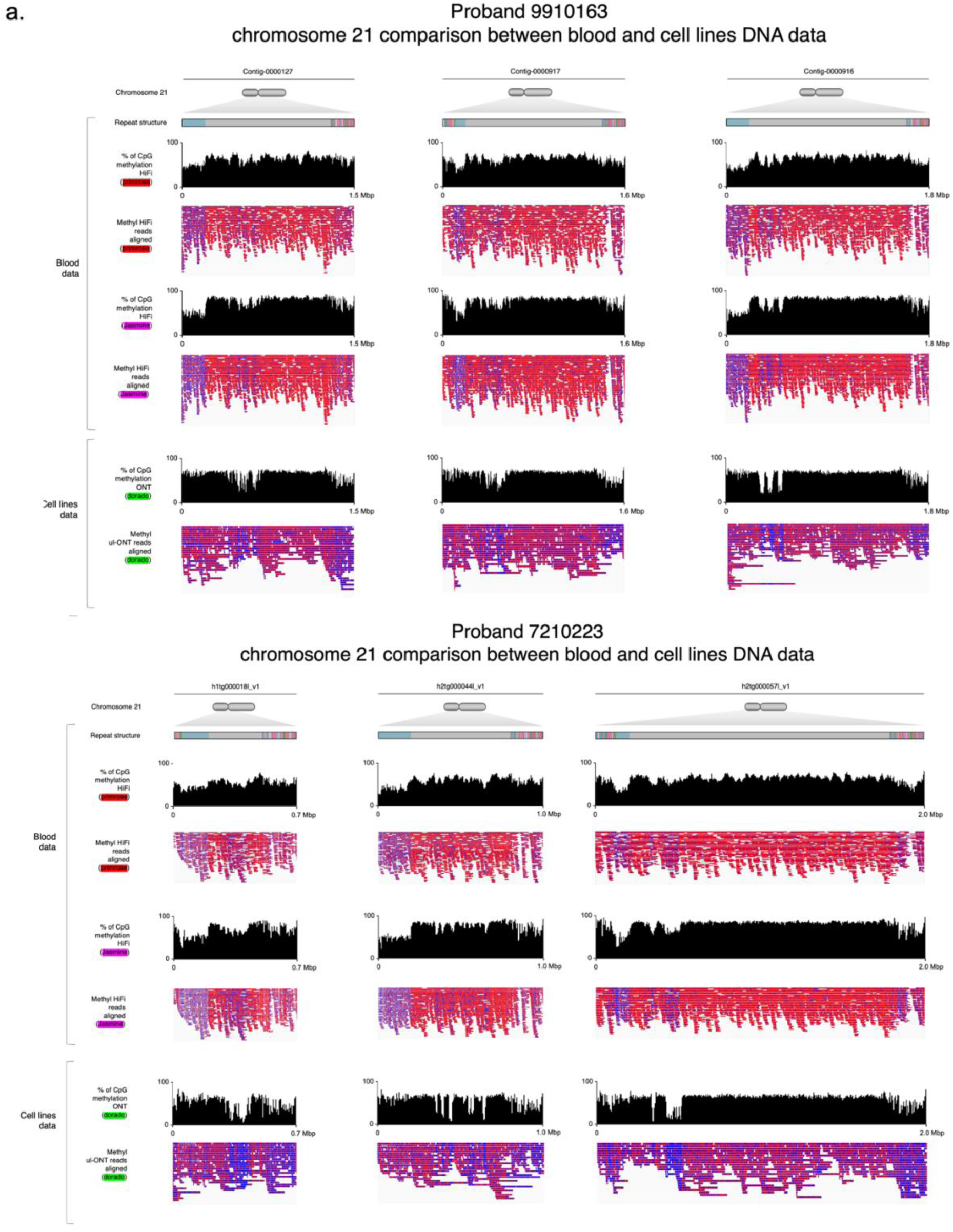
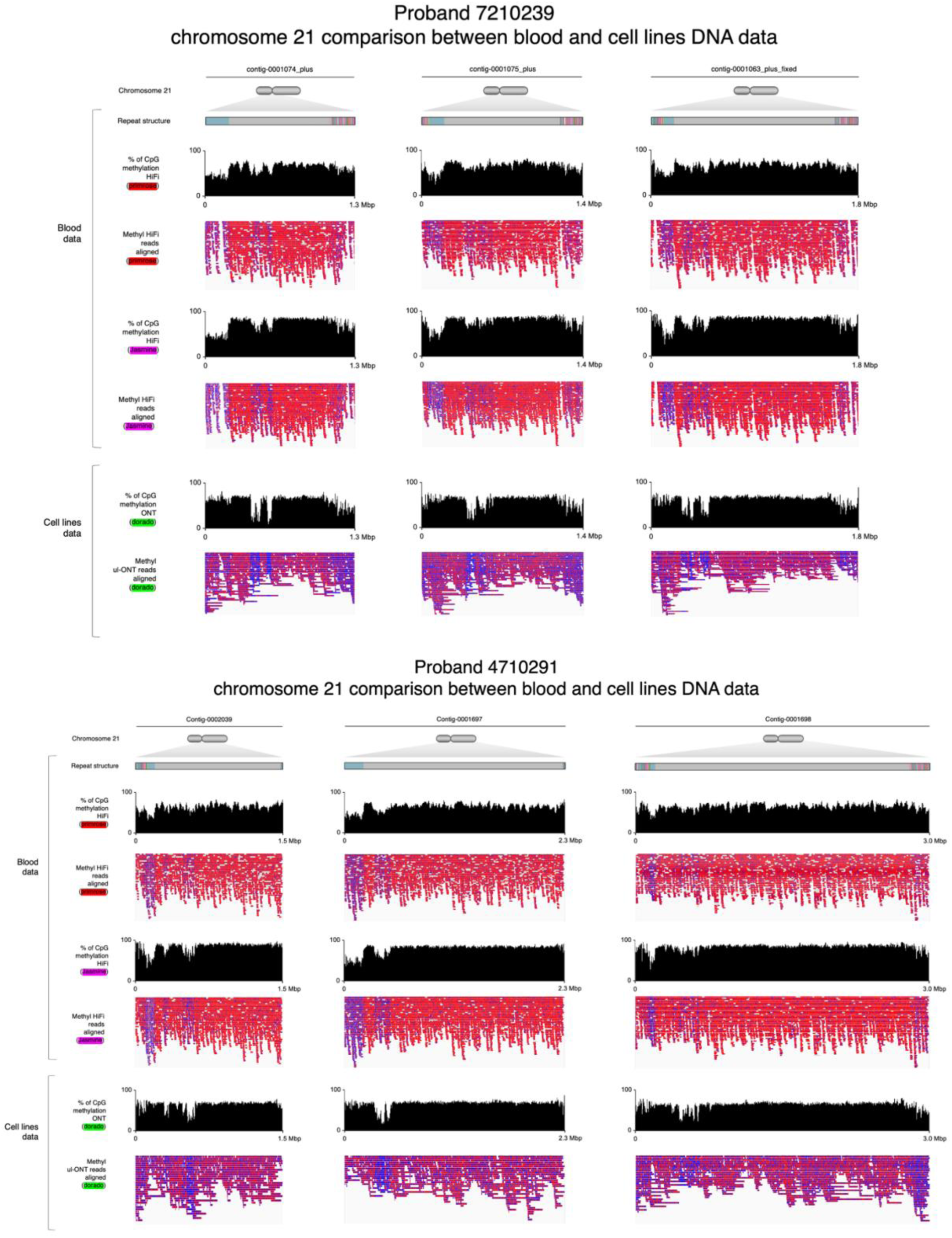
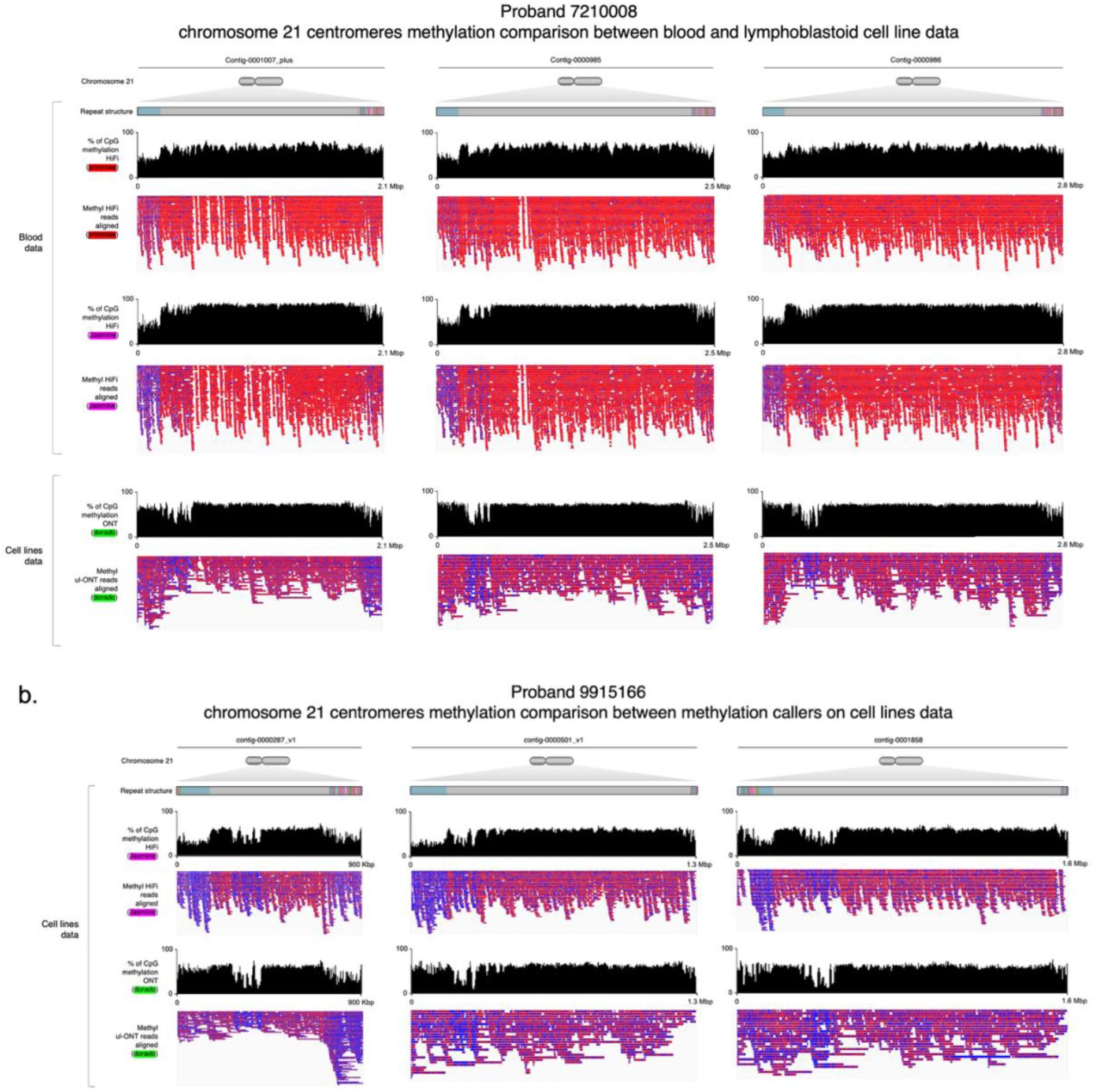
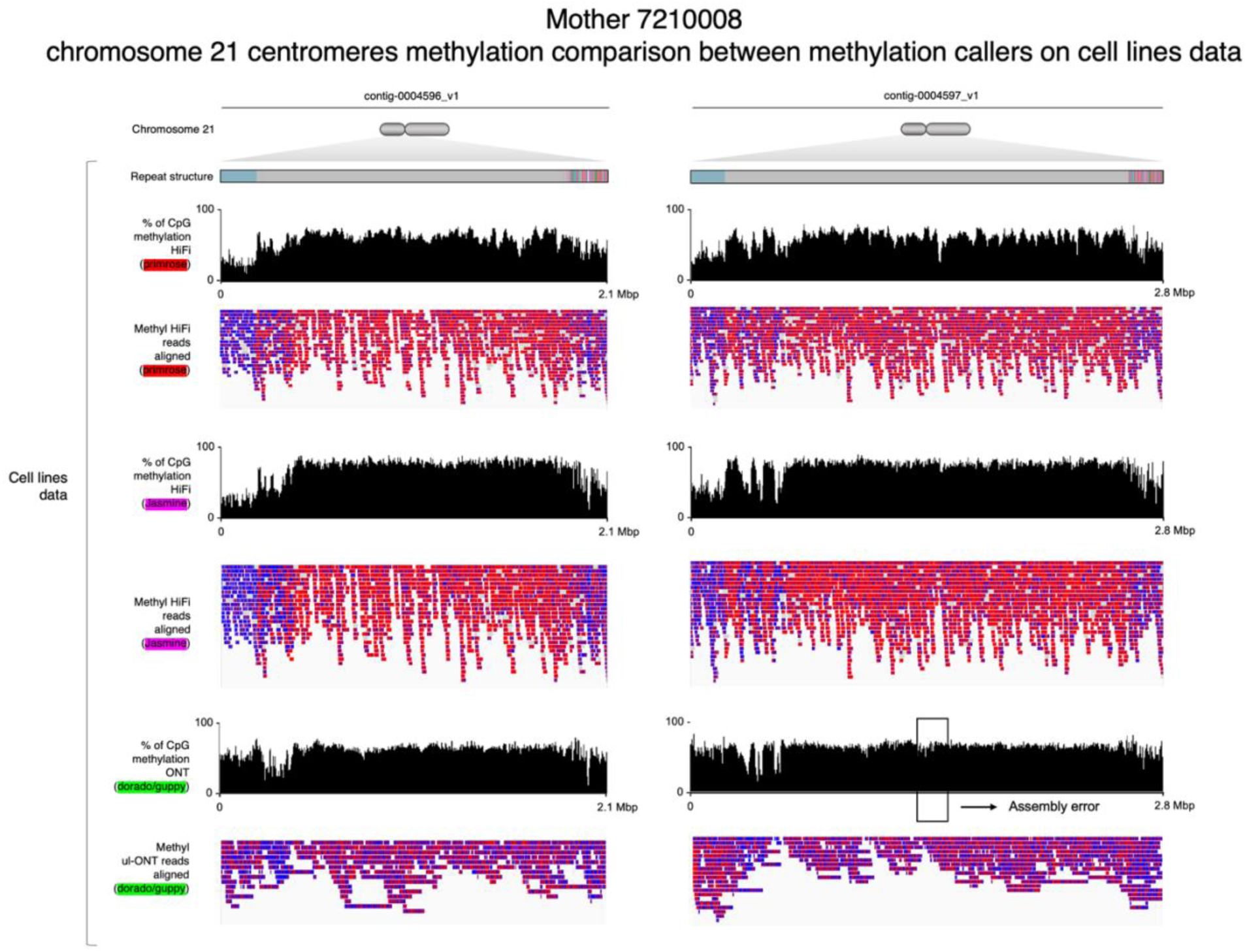
Comparison of methylation profiles of chr21 centromeres in blood and cell line data. **a)** Chr21 centromere methylation profiles detected from HiFi (blood DNA) and UL-ONT (lymphoblastoid cell line DNA) in probands 9910163.00, 7210223.00, 7210239.00, 4710291.00, and 7210008.00. **b)** Performance comparison of methylation callers on cell line data in proband 9915166.00 and mother 7210008.20.

**Supplementary Figure 9.**
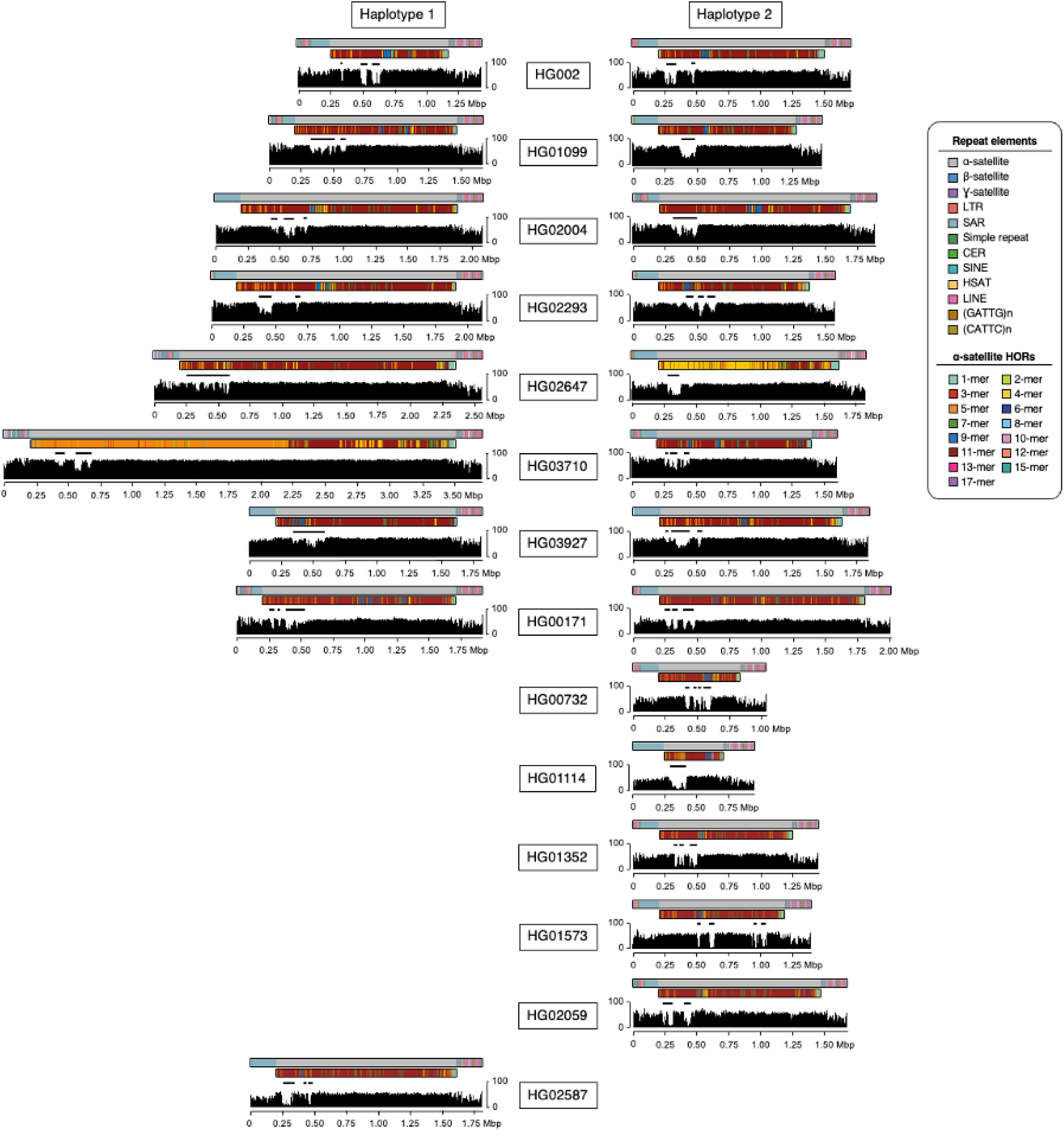
Extended version of Figure 3c representing the methylation profiles and CDR positions in 22 population chr21 centromere haplotypes.

**Supplementary Figure 10.**
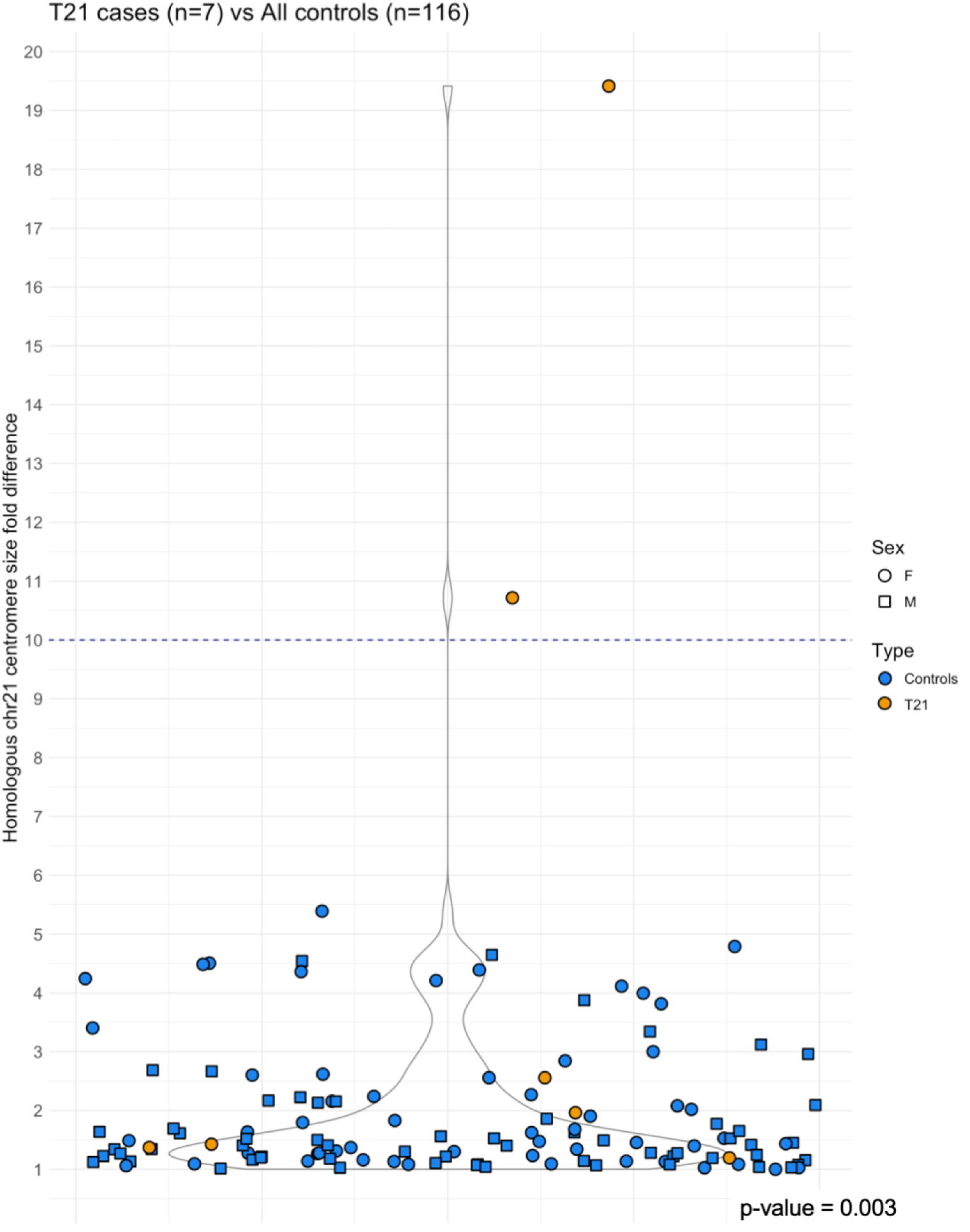
Homologous chr21 centromere size fold differences in T21 centromeres (only maternally inherited) and population samples. T21 (orange) and population (blue) samples are represented with different colors; sex is represented with different shapes. P-value indicates a significant difference in asymmetry cases between the two groups.

---

**Supplementary Table 1.** All HOR monomers identified in 35 highly curated population chr21 centromeres.

**Supplementary Table 2.** Information on families with T21.

**Supplementary Table 3.** All chr21 centromeres described in the manuscript.

## LIST OF ABBREVIATIONS

CDR: centromere dip region
chr21: chromosome 21
FISH: fluorescence *in situ* hybridization
Gbp: Gigabase pairs
IF: immunofluorescence
kbp: kilobase pairs
kya: thousand years ago
H1/2/3/4: haplotype 1/2/3/4
HiFi: high fidelity
HGSVC: Human Genome Structural Variation Consortium
HPRC: Human Pangenome Reference Consortium
HOR: higher-order repeat
MMI/IIE: maternal meiosis I/II error
Mbp: Megabase pairs
PacBio: Pacific Biosciences
T21: Trisomy 21
T2T: telomere-to-telomere
UL-ONT: ultra-long Oxford Nanopore Technologies sequencing

